# Molecular evidence for sweeping discontinuity between peracarid (Crustacea) fauna of Macaronesian islands and nearby continental coasts: over fifty candidate endemic species

**DOI:** 10.1101/2021.06.22.449383

**Authors:** Pedro E Vieira, Andrea Desiderato, Sofia L Azevedo, Patricia Esquete, Filipe O Costa, Henrique Queiroga

## Abstract

Oceanic islands are recognized evolutionary hotspots for terrestrial organisms, but little is known about their impact on marine organisms’ evolution and biogeography. The volcanic archipelagos of Macaronesia occupy a vast and complex region which is particularly suitable to investigate marine island biogeography.

In this study, we used mitochondrial DNA sequences to investigate the genetic diferentiation between the populations from Webbnesia (i.e. Madeira, Selvagens and Canaries) and adjacent coasts, of 23 intertidal peracarid species. All species had unexpectedly high intraspecific genetic distances, reaching more than 20% in some cases. Between 79 and 95 Molecular Operational Taxonomic Units (MOTUs) were found in these species. Webbnesia populations displayed an impressive genetic diversity and high endemicity, with 83% of the MOTUs being private to these islands, particularly La Palma and Madeira. Network analyses suggested higher similarity between Webbnesia and Azores than with adjacent continental coasts.

These results reveal an unanticipated and sweeping biogeographic discontinuity of peracaridean fauna between Webbnesia and the Iberian Peninsula, raising suspicion about the possible occurrence of identical patterns in other groups of marine invertebrates in the region. We emphasize the unique genetic heritage hosted by these islands, underlining the need to consider the fine scale endemicity in marine conservation efforts.

## Introduction

The marine realm is generally considered to have lower habitat diversity and higher connectivity than terrestrial habitats [1,2]. Non-marine biota inabithing oceanic islands have to cross the ocean to disperse and are more prone to isolation than marine organisms [3]. However, several studies have been indicating an increasing number of discountinuities between and within marine bioregions, possibly driven by constraints in dispersal and gene flow, that only recently started to be noticed and reported (e.g. [4,5]). Moreover, it is known that even geographically close islands [6–8] may comprise distinct marine coastal communities in response to local biotic and abiotic factors.

Recent studies on the marine biota of Macaronesia, sustain that this group of 31 islands belonging to five archipelagos (i.e. Azores, Madeira, Selvagens, Canaries, Cape Verde) in the Northeast Atlantic (NEA), comprise in fact not one, but three distinct bioregions. For example, Cabo Verde differs significantly from the other Macaronesian archipelagos and appears to be a subprovince within the West African Transition province [9–11], while the remaining archipelagos may belong to the Lusitanian province [11,12]. Because Madeira, Selvagens and Canaries share a higher affinity in their biota, it was proposed that these archipelagos should be grouped in a separate ecoregion named “Webbnesia”, leaving the Azores as an independent ecoregion by itself [11].

Recently, with the support of molecular tools, we have found cryptic diversity within the isopod *Dynamene edwardsi* (Lucas, 1849) [13] and in the amphipod family Hyalidae [14] occurring in Macaronesia. Our studies suggested segregation among islands and a possible discontinuity between Webbnesian fauna and the adjacent continental landmasses.

Peracarids are abundant benthic crustaceans in marine coasts that have presumably lower dispersal capacities due to lacking planktonic larvae, thereby being particularly suited to investigate biogeographic discontinuities in the open ocean. In this study, we aimed to use the cytochrome c oxidase subunit I (COI) DNA barcoding region [15] to conduct a comprehensive parallel screening of genetic differentiation across populations from the NEA of 23 morphospecies of Amphipoda, Isopoda and Tanaidacea. In particular, we aim to probe the occurrence of cryptic diversity by investigating the suspected genetic and taxonomic discountinuities between the above-mentioned presumptive bioregions.

## Material and Methods

Peracarid specimens were collected in the archipelagos of Azores, Madeira, Selvagens and Canaries and in the continental coasts of Morocco and Iberian Peninsula (figure 1). Morphology-based taxonomic identification was performed consulting specialized literature. Sampling details and literature used can be accessed in the supplementary material.

**Figure 1.**
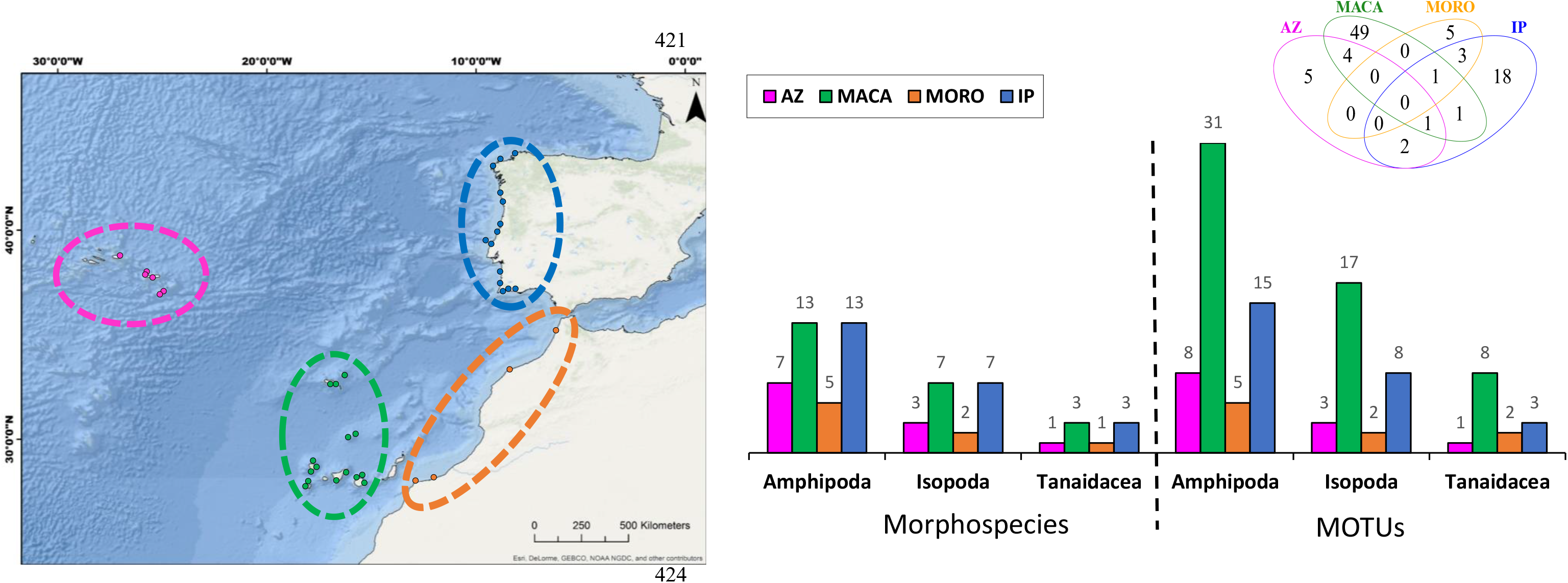
Sampling locations (left) and number of morphospecies and MOTUs retrieved for each region for each order (right). The Venn diagram shows the total number of endemic and shared MOTUs between regions. Number of consensus MOTUs accordingly with Table 2. Co-ordinates can be consulted in Table S1. For interactive map, see http://rpubs.com/Vieira/PeracaridaNE. In the interactive map, in the right-top corner, it is possible to choose the different species and verify the sampling locations for each species. The records of *Stenothoe monoculoides* from North Sea are only displayed in the interative map. The interactive map was created with the package “leaflet” [65], through the software R 3.5.0 [21]. Az - Azores; MACA - Webbnesia; MORO - Morocco; IP - Iberian Peninsula.

According to the main hypothesys, specimens of each species were chosen from two main regions (Iberian Peninsula and Webbnesia), following the genetic differentiation observed between these regions in previous works [13,14]. The first group included the specimens sampled in Iberian Peninsula (IP) and the second included the specimens collected in Webbnesia, i.e. Madeira, Selvagens and Canaries archipelagos (MACA). Only the sequences of *Stenothoe monoculoides* (Montagu, 1813) were from the North Sea, because there were no public data available from the Iberian Peninsula. However, our unpublished data derived from metabarcoding already detected this species in Northwest of Spain and was confirmed as the same haplotype as the one from North Sea. Therefore, we are confident that this morphospecies occurs in Iberian Peninsula. In addition, when present, specimens from Morocco (MORO) and Azores (AZ), were added to the main experimental design (figure 1; supplementary material, Table S1).

DNA extraction, COI amplification, PCR products purification and sequencing were performed for each specimen following [14]. The other sequences were obtained in our previous works [13,14,16,17] and from [18] (see supplementary material, Tables S1-S2, for list of primers, number of specimens in each species and source). A common fragment of 520 base pair was obtained and used in subsequent analyses. Maximum and mean pairwise distances (p-distances) for COI within each morphospecies were calculated in general and within groups in MEGA 7.0 [19].

To assess the presence of cryptic species (i.e. multiple molecular operational taxonomic units - MOTUs) in each morphospecies [20], five methods were applied to the dataset: automatic barcode gap analysis (ABGD), BOLD (BINs), bayesian Poisson Tree Partition (bPTP), Generalized Mixed Yule Coalescent (GMYC) and TCS (details can be consulted in the supplementary material). A majority rule (i.e. most commom number of MOTUs for each species) was applied and, in case of a tie, a conservative approach was applied choosing the lowest number of MOTUs.

Chord diagrams were built in R 3.5.0 [21] with the package ‘chorddiag’ [22] to inspect the number of MOTUs endemic to each island (including Iberia and Morocco) and region (i.e. MACA, IP, AZ, MORO), and amount of shared ones. Community detection representations (based on shared and private MOTUs between/within locations) were calculated with the R packages ‘igraph’ [23] and ‘visNetwork’ [24].

## Results

### Molecular analyses and MOTUs delimitation

A total of 483 sequences were analysed, of which 173 were produced in this study, belonging to 23 morphospecies. Mean intraspecific distance (ISD) varied between 1.81% (*Ampithoe ramondi* Audouin, 1826) and 17.16% (*Janira maculosa* Leach, 1814), while Maximum ISD was higher than 3% for all species (Table 1). Mean p-distances between IP and MACA regions were always higher than 3%, with the highest value observed in the isopod *Anthura gracilis* (Montagu, 1808) (28%, Table 1).

**Table 1.** Presence (*●*) of the peracaridean species used in this study in each region. Mean and Maximum (Max) intraspecific distance (ISD) for each species. The Mean p-distance between the Iberian Peninsula (IP) and Webbnesia (MACA) populations for each morphospecies is also displayed. AZ – Azores; MORO – Morocco.

| Order | Species | MACA | IP | AZ | MORO | Mean ISD | Max ISD | Mean p-distances between IP and MACA |
| --- | --- | --- | --- | --- | --- | --- | --- | --- |
| Amphipoda | *Ampithoe helleri* | ● | ● |  |  | 0.0715 | 0.1327 | 0.1230 |
|  | *Ampithoe ramondi* | ● | ● | ● |  | 0.0181 | 0.0385 | 0.0341 |
|  | *Ampithoe riedli* | ● | ● |  | ● | 0.0439 | 0.0827 | 0.0782 |
|  | *Apohyale perieri* | ● | ● | ● |  | 0.0483 | 0.1135 | 0.0770 |
|  | *Apohyale stebbingi* | ● | ● | ● | ● | 0.1243 | 0.2000 | 0.1574 |
|  | *Caprella acanthifera* | ● | ● | ● | ● | 0.0805 | 0.1462 | 0.1374 |
|  | *Elasmopus pecteniscus* | ● | ● |  | ● | 0.0381 | 0.0635 | 0.0583 |
|  | *Jassa herdmanni* | ● | ● | ● |  | 0.0751 | 0.1362 | 0.1237 |
|  | *Podocerus variegatus* | ● | ● |  |  | 0.0613 | 0.1019 | 0.0974 |
|  | *Protohyale schmidtii* | ● | ● | ● | ● | 0.0693 | 0.1346 | 0.1087 |
|  | *Quadrinemaera inaequipes* | ● | ● |  |  | 0.0911 | 0.1596 | 0.1357 |
|  | *Serejohyale spinidactylus* | ● | ● | ● |  | 0.1152 | 0.1769 | 0.1348 |
|  | *Stenothoe monoculoides* * | ● | ● |  |  | 0.1637 | 0.2765 | 0.2765 |
| Isopoda | *Anthura gracilis* | ● | ● | ● | ● | 0.1521 | 0.2846 | 0.2800 |
|  | *Campecopea lusitanica* | ● | ● |  |  | 0.1012 | 0.1981 | 0.1226 |
|  | *Cymodoce truncata* | ● | ● | ● |  | 0.1263 | 0.2019 | 0.1619 |
|  | *Dynamene edwardsi* | ● | ● | ● | ● | 0.1140 | 0.1865 | 0.1643 |
|  | *Gnathia maxillaris* | ● | ● |  |  | 0.1324 | 0.2038 | 0.2000 |
|  | *Janira maculosa* | ● | ● |  |  | 0.1715 | 0.2673 | 0.2564 |
|  | *Joeropsis brevicornis* | ● | ● |  |  | 0.1252 | 0.2500 | 0.2462 |
| Tanaidacea | *Apseudopsis latreilii* | ● | ● |  |  | 0.1674 | 0.2404 | 0.2372 |
|  | *Tanais dulongii* | ● | ● |  | ● | 0.0840 | 0.1192 | 0.1150 |
|  | *Tanais grimaldii* | ● | ● | ● |  | 0.0919 | 0.1481 | 0.1065 |

\**Stenothoe monoculoides* was retrieved from North Sea instead of IP.

**Table 2.**
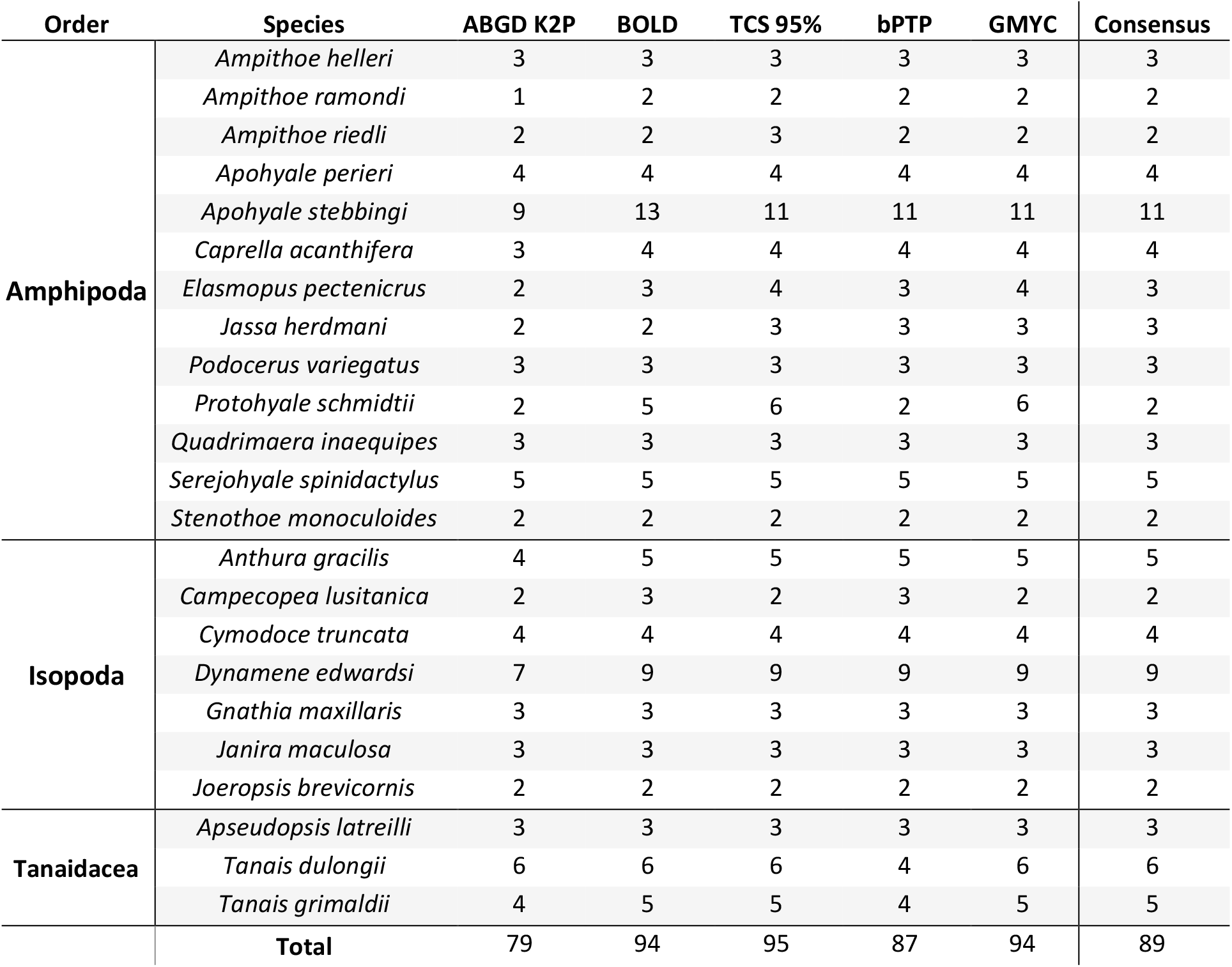
Number of MOTUs accordingly to different molecular species delineation methods for each morphospecies.

| Order | Species | ABGD K2P | BOLD | TCS 95% | bPTP | GMYC | Consensus |
| --- | --- | --- | --- | --- | --- | --- | --- |
| Amphipoda | *Ampithoe helleri* | 3 | 3 | 3 | 3 | 3 | 3 |
|  | *Ampithoe ramondi* | 1 | 2 | 2 | 2 | 2 | 2 |
|  | *Ampithoe riedli* | 2 | 2 | 3 | 2 | 2 | 2 |
|  | *Apohyale perieri* | 4 | 4 | 4 | 4 | 4 | 4 |
|  | *Apohyale stebbingi* | 9 | 13 | 11 | 11 | 11 | 11 |
|  | *Caprella acanthifera* | 3 | 4 | 4 | 4 | 4 | 4 |
|  | *Elasmopus pecteniscus* | 2 | 3 | 4 | 3 | 4 | 3 |
|  | *Jassa herdmani* | 2 | 2 | 3 | 3 | 3 | 3 |
|  | *Podocerus variegatus* | 3 | 3 | 3 | 3 | 3 | 3 |
|  | *Protohyale schmidtii* | 2 | 5 | 6 | 2 | 6 | 2 |
|  | *Quadrinemaera inaequipis* | 3 | 3 | 3 | 3 | 3 | 3 |
|  | *Serejohyale spinidactylus* | 5 | 5 | 5 | 5 | 5 | 5 |
|  | *Stenothoe monoculoides* | 2 | 2 | 2 | 2 | 2 | 2 |
| Isopoda | *Anthura gracilis* | 4 | 5 | 5 | 5 | 5 | 5 |
|  | *Campecopea lusitanica* | 2 | 3 | 2 | 3 | 2 | 2 |
|  | *Cymodoce truncata* | 4 | 4 | 4 | 4 | 4 | 4 |
|  | *Dynamene edwardsi* | 7 | 9 | 9 | 9 | 9 | 9 |
|  | *Gnathia maxillaris* | 3 | 3 | 3 | 3 | 3 | 3 |
|  | *Janira maculosa* | 3 | 3 | 3 | 3 | 3 | 3 |
|  | *Joeropsis brevicornis* | 2 | 2 | 2 | 2 | 2 | 2 |
| Tanaidacea | *Apseudopsis latreilli* | 3 | 3 | 3 | 3 | 3 | 3 |
|  | *Tanais dulongii* | 6 | 6 | 6 | 4 | 6 | 6 |
|  | *Tanais grimaldii* | 4 | 5 | 5 | 4 | 5 | 5 |
|  | **Total** | 79 | 94 | 95 | 87 | 94 | 89 |

The molecular species delimitation methods retrieved between 79 (ABGD) and 95 (TCS) MOTUs (Table 2). Between 41 and 53 MOTUs were present among the 13 amphipods, between 25 and 29 MOTUs in the seven isopods and between 11 and 14 MOTUs in the three tanaidaceans (Table 2, supplementary material Figs. S1-3). The consensus number of MOTUs was 89 (Table 2), with a minimum of two in six morphospecies and maximum of 11 in *Apohyale stebbingi* Chevreux, 1888 (Table 2, supplementary material, Fig. S4).

### Peracarid community analysis

The MACA region harboured more MOTUs than the IP region (56 and 26 respectively; figure 1, supplementary material, Fig. S4), with the islands of La Palma (19), Madeira (17) and Gran Canaria (14) with the highest number of MOTUs. No more than four MOTUs were shared between islands and only three were shared between MACA and IP (figure 1). La Palma and Madeira were the islands with the highest number of private MOTUs (12 and 8 respectively), with MACA displaying 49 endemic MOTUs and IP only 18 (figure 1; supplementary material, Fig. S4).

The artificial networks of the islands including Morocco and Iberian Peninsula, retrieved Multilevel (modularity:0.080; figure 2A), Spinglass (modularity:0.110; figure 2B), Edge betweenness (modularity:0.020; figure 2C) and Walktrap (modularity:0.023; figure 2D) as the most fitting community detection algorithms to our data. All these algorithms grouped Canaries and Azores together, with Madeira and Selvagens showing different patterns (depending on the algorithm), and Morocco and IP in separate clusters. When regions were used, the network retrieved Multilevel and Spinglass (modularity: 0.061; figure 2E) algorithms. Both retrieved the same topology (figure 2E), with pairs MORO-IP and AZ-MACA clustering together.

**Figure 2.**
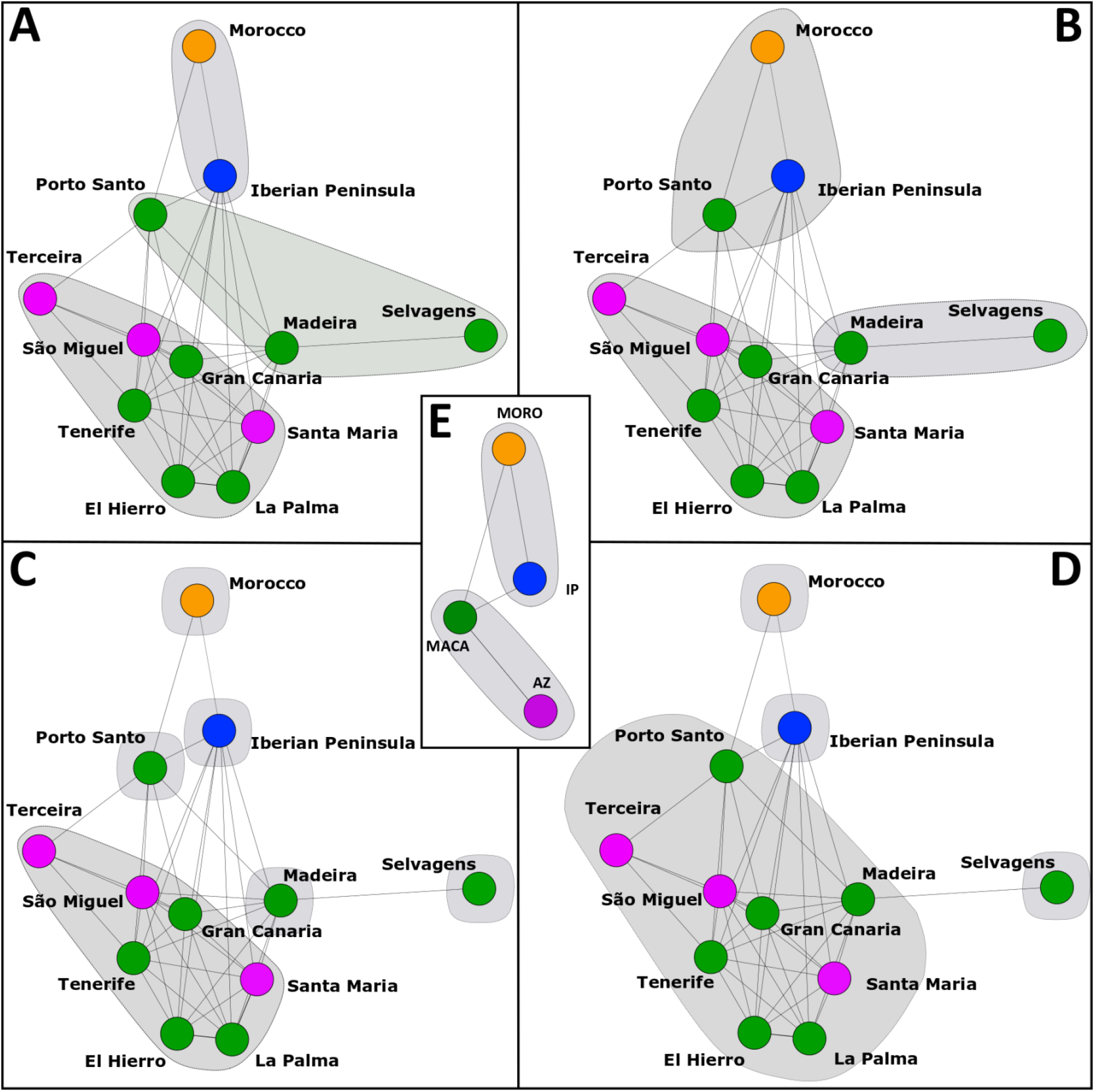
Network scenarios resulted from the best algoritms tested using locations (A-D) and regions (E). A: multi level; B: spinglass; C: edge betweenness; D: walktrap; E: multi level and spinglass. Interactive networks can be accessed at https://rpubs.com/Vieira/Peracaridab and at https://rpubs.com/Vieira/Peracaridab. Az - Azores; MACA - Webbnesia; MORO - Morocco; IP - Iberian Peninsula.

## Discussion

Compared to terrestrial fauna, very little is known about the biogeography and evolution of insular marine fauna [25]. The common perception for Macaronesia’s marine invertebrate fauna is that many species are shared with mainland coasts of NW Africa and Iberia, hence a basal faunistic continuity is assumed. The absence of any obvious geographic barriers for marine organisms’ dispersal in the region intituively reinforces this perception. Our findings appear to contradict this view, at least in what concerns peracarids. We have found: i) extensive and profound genetic differentiation between peracarid populations from Iberia and Webbnesia; ii) extensive peracarid endemic diversity in Webbnesia, patent in 48 well-supported and highly divergent MOTUs; and iii) geographic segregation among Webbnesia’s MOTUs, including many private to only one or a few islands. Our study captured for the first time this faunistic discontinuity because, to this date, it is probably the most extensive marine invertebrate metaspecies screening of genetic differentiation between Atlantic continental and islands populations.

### Diversification and evolution of peracarids and other marine invertebrates in Macaronesia

The 23 species here examined displayed completely sorted MOTUs between Iberia and Webbnesia, which were well supported by multiple clustering methods. The large amount of COI data available for animals indicates that COI-based MOTUs commonly correspond to separate species [26,27]. The genetic distances within morphospecies we observed are above the intrapecific range reported in comprehensive studies with crustaceans [17, 18, 28]. Even considering the top range of COI evolutionary rates estimated for crustaceans [29], these distances indicate long-term evolutionary divergence and suggest that these may be separate species.

Phylogeographic discontinuities have been reported in marine environments worldwide, e.g. [5, 30–34], as the notorious case of the Wallace’s line in the Makassar Strait [4], but little is known for the NEA. Differentiation between populations of Webbnesia and those from Iberian Peninsula was also found in taxa with a planktonic phase such as sponges [35], molluscs [36,37] and fish [38], suggesting a phylogeographic discontinuity for marine fauna in general, despite the potential for larval transport by recurring oceanographic features. Long-distance dispersal in peracarideans is mainly due to stochastic events through rafting on floating objects or mediated by human vectors [39,40]. Notwithstanding the different lifestyles of each peracaridean species, a deep genetic differentiation was transversal between the populations from Webbnesia and adjacent continental coasts, suggesting that other factors such as "post-colonization monopolization” or the islands caracteristics (e.g. [41–44]) may play a major role in the geographic seggregation of these species. Moreover, considering the deep genetic divergences found, it is probable that the populations’ differentiation preceed the last glaciation maximum [45] and occurred thousands to millions of years ago [13].

While a clear differentiation between Webbnesia and Iberian populations was patent in all the examined species, the populations from Azores and Morocco displayed specificities depending on the taxon. The biota of Azores and Webbnesia is usually presumed similar [46], due to the currents’ patterns during interglacial periods [11, 47]. Previous studies showed genetic similarities between marine invertebrates of these archipelagos [35,48], while others suggest stronger affinities between Azorean and Iberian populations [36]. In this study, both patterns were observed, although with higher support for the Azores-Webbnesia connection than for the Azores-Iberian Peninsula. Moreover, our data suggests a higher genetic proximity between Moroccan and Iberian populations, contradicting other studies that relate the populations from Morocco with those from Macaronesia due to their vicinity [32, 46, 49, 50].

During glacial periods, the isolation of Webbnesia’s islands may have been significantly reduced when compared to present geographic distances [51], due to a lower sea level, greater surface area and exposure of the currently submerged islands that could have served as stepping-stones. This factor, together with the similarity of habitats and geographic proximity between Madeira, Selvagens and Canaries, may explain the high number of shared MOTUs within Webbnesia (figure 2).

### Implications for marine biodiversity conservation and management

Species are commonly used as framework for conservation strategies since they constitute the basic units for distributional and habitat studies in biodiversity assessments. However, with the emergence of molecular methods, the importance of molecular evidence for species delineation [52] arose as a critical contribution to understand evolution and inform conservation strategies. Using molecular methods, MOTUs could be considered as the functional units of biodiversity and might act as proxies for estimating diversity [53]. Concepts such as “Evolutionary Significant Units” (ESU) may help surpass the limitations imposed by rigid species boundaries [54], enabling the recognition of pertinent infraspecific units for the purpose of biodiversity conservation [55] and connectivity [56]. Hence, regardless of the formal species boundaries of the peracarids here investigated, it appears there is at least an extraordinary level of endemicity of genetic lineages with very small ranges, frequentely no larger than the island that harbours them.

The preservation of genetic diversity is an essential factor in the design of marine conservation areas which should therefore include domains that incorportate fundamental evolutionary processes [57]. In the marine environment, priority should be given to the conservation of those species most vulnerable to human activities and those with populations dangerously affected. Due to their small size and isolation, island and endemic species are more likely to extinguish than continental or non-endemic species [58]. Moreover, the human activities that mainly affect the marine environment usually take place in coastal areas, whose extension is limited and where the highest marine productivity is reached [57,59].

Marine invertebrates are rarely contemplated in marine protected areas, despite benefitting greatly from these programs [60], and little information is available about the status of each species/MOTU/ESU/population in Macaronesia. Remarkably, neither Webbnesia or the Azores were included yet in the “Ecologically or Biologically Significant Marine Areas” (EBSAs; www.cbd.int/ebsa/ebsas) [61]. However, both would probably qualify if there was a wider awareness of their unique endemic marine diversity and vulnerability. Indeed, the extent of the taxonomic and genetic diversity harboured by the marine invertebrates from Macaronesia is still poorly known [11,62] and there is an urgent need to accelerate its inventory, particularly using molecular tools, given that at current rates it will take decades till completion [63]. To protect, manage and conserve the unique biological heritage of these archipleagos, it is crucial that the fine-scale endemicity of marine organisms is considered in the design of more effective networks of marine protected areas.

### Conclusions

This study provides compelling evidence for a sweeping discontinuity in shallow-water peracarid fauna between Webbnesia and nearby continental coasts. We also found rampant endemic peracarid diversification in these archipelagos, and multiple cases of clear geographic sorting of MOTUs even among islands separated by no more than 60 km. These findings challenge the intuitive perception of faunistic continuity of marine organisms between islands and nearby mainland, somewhat downplaying the role of contemporary dispersal and connectivity as a main explanation for the biogeography of insular marine organisms. Indeed, founder effects, mechanisms of monopolization and preemptive exclusion [13, 41, 64], coupled with the islands’ configuration [44], may have a more prevalent role in the elucidation of contemporary biogeographies of islands’ shallow water invertebrates than previously acknowledged. We hope these results may rise the awareness on the need of considering a larger variety of taxa for the identification of protected areas shedding light into the poorly known island biogeography of marine organisms.

## Acknowledgements

The authors wish to thank the colleagues who helped during fieldwork, sample processing and/or laboratory work: Tavares M and Santos R (University of Algarve, Portugal), Ladeiro B, Peteiro L, Gomes I, Albuquerque R, Guimarães B and Fuente N (University of Aveiro, Portugal) and Gomes N (University of Minho, Portugal). Aditionally, thanks to Carvalho D in name of the Portuguese Museum of Natural History and Science of Lisbon for supplying material from the EMEPC/M@rBis/Selvagens2010 and EMAM/PEPC_M@rBis/2011 campaigns to Selvagens. Thanks to Bellisario B for feedback regarding network analysis. Finally, thanks to Ferreira EL for the use of some equipments.

This work was supported by the project “DiverseShores - Testing associations between genetic and community diversity in European rocky shore environments (PTDC/BIA-BIC/114526/2009)” funded by the Fundação para a Ciência e Tecnologia (FCT) under the COMPETE programme supported by the European Regional Development Fund. FCT also supported a PhD grant to PEV (SFRH/BD/86536/2012). Thanks to FCT/MCTES are also due for the financial support to CESAM (UIDP/50017/2020+UIDB/50017/2020), through national funds. PE was funded through FCT in the scope of the framework contract foreseen in the numbers 4, 5 and 6 of the article 23 of the Decree-Law 57/2016, of August 29, changed by Law 57/2017, of July 19.

## Data Accessibility

All new DNA sequences generated in this work were deposited in BOLD under the projects (PMACA: “Peracarida Macaronesia vs IberiaPeninsula” and PERAC: “Peracarida New data”). All the data used in this work is available in the BOLD dataset DS-PMACA: “Peracarida Macaronesia vs IberiaPeninsula”. All R scripts are available at https://github.com/pedroemanuelvieira/Macaronesiadiscontinuity.

## Author Contributions

PEV, FOC and HQ designed the research plan; PEV, AD and SLA performed the research and analysed the data; PEV, PE and AD identified the specimens; PEV wrote the original manuscript; all the authors contributed with suggestions, to the manuscript structure and reviewed the manuscript final version.

## References

1. Gray JS. 1997 Marine biodiversity: patterns, threats and conservation needs. Biodivers. Conserv. 6, 153–175.

2. Appeltans W et al. 2012 The Magnitude of Global Marine Species Diversity. Curr. Biol. 22, 2189–2202.

3. Cox SC, Carranza S, Brown RP. 2010 Divergence times and colonization of the Canary Islands by Gallotia lizards. Mol. Phylogenet. Evol. 56, 747–757. (doi: 10.1016/j.ympev.2010.03.020).

4. Barber PH, Palumbi SR, Erdmann MV, Moosa MK. 2000 A marine Wallace’s line? Nature 406, 692–693.

5. Bellisario B, Camisa F, Abbattista C, Cimmaruta R. 2019 A network approach to identify bioregions in the distribution of Mediterranean amphipods associated with Posidonia oceanica meadows. PeerJ 7, e6786. (doi: 10.7717/peerj.6786).

6. Fernández-Palacios JM et al. 2016 Towards a glacial-sensitive model of island biogeography. Glob. Ecol. Biogeogr. 25, 817–830. (doi: 10.1111/geb.12320).

7. Rijsdijk KF et al. 2014 Quantifying surface-area changes of volcanic islands driven by Pleistocene sea-level cycles: biogeographical implications for the Macaronesian archipelagos. J. Biogeogr. 41, 1242–1254. (doi: 10.1111/jbi.12336).

8. Ávila SP et al. 2018 Global change impacts on large-scale biogeographic patterns of marine organisms on Atlantic oceanic islands. Mar. Pollut. Bull. 126, 101–112. (doi: 10.1016/j.marpolbul.2017.10.087).

9. Spalding MD et al. 2007 Marine Ecoregions of the World: A Bioregionalization of Coastal and Shelf Areas. BioScience 57, 573–583. (doi: 10.1641/B570707).

10. Tuya F, Haroun RJ. 2009 Phytogeography of Lusitanian Macaronesia: biogeographic affinities in species richness and assemblage composition. Eur. J. Phycol. 44, 405–413. (doi: 10.1080/09670260902836246).

11. Freitas R et al. 2019 Restructuring of the ‘Macaronesia’ biogeographic unit: A marine multi-taxon biogeographical approach. Sci. Rep. 9, 15792. (doi: 10.1038/s41598-019-51786-6).

12. Almada VC et al. 2013 Complex origins of the Lusitania biogeographic province and northeastern Atlantic fishes. Front. Biogeogr. 5, 1. (doi: 10.21425/F5FBG14493).

13. Vieira PE et al. 2019 Deep segregation in the open ocean: Macaronesia as an evolutionary hotspot for low dispersal marine invertebrates. Mol. Ecol. 28, 1784–1800. (doi: 10.1111/mec.15052).

14. Desiderato A et al. 2019 Macaronesian islands as promoters of diversification in amphipods: The remarkable case of the family Hyalidae (Crustacea, Amphipoda). Zool. Scr. 48, 359–375. (doi: 10.1111/zsc.12339).

15. Hebert PDN, Cywinska A, Ball SL, deWaard JR. 2003 Biological identifications through DNA barcodes. Proc. R. Soc. Lond. B Biol. Sci. 270, 313–321. (doi: 10.1098/rspb.2002.2218).

16. Lobo J et al. 2013 Enhanced primers for amplification of DNA barcodes from a broad range of marine metazoans. BMC Ecol. 13, 34. (doi: 10.1186/1472-6785-13-34).

17. Lobo J et al. 2016 Contrasting morphological and DNA barcode-suggested species boundaries among shallow-water amphipod fauna from the southern European Atlantic coast. Genome (doi:10.1139/gen-2016-0009).

18. Raupach MJ et al. 2015 The Application of DNA Barcodes for the Identification of Marine Crustaceans from the North Sea and Adjacent Regions. PLOS ONE 10, e0139421. (doi: 10.1371/journal.pone.0139421).

19. Kumar S, Stecher G, Tamura K. 2016 MEGA7: Molecular Evolutionary Genetics Analysis Version 7.0 for Bigger Datasets. Mol. Biol. Evol. 33, 1870–1874. (doi: 10.1093/molbev/msw054).

20. Fišer C, Robinson CT, Malard F. 2018 Cryptic species as a window into the paradigm shift of the species concept. Mol. Ecol. 27, 613–635. (doi: 10.1111/mec.14486).

21. R Core Team. 2018 R: A language and environment for statistical computing. R Foundation for Statistical Computing, Vienna, Austria. https://www.R-project.org/.

22. Flor M. 2016 chorddiag: Interactive Chord Diagrams. R package version 0.1.2. http://github.com/mattflor/chorddiag/.

23. Csardi G, Nepusz T. 2006 The igraph software package for complex network research. InterJournal Complex Systems. http://igraph.org.

24. Almende B, Thieurmel B, Robert T. 2018 visNetwork: Network Visualization using ‘vis.js’ Library. R package version 2.0.4. https://CRAN.R-project.org/package=visNetwork.

25. Dawson MN. 2016 Island and island-like marine environments. Glob. Ecol. Biogeogr. 25, 831–846. (doi: 10.1111/geb.12314).

26. Teixeira MAL. et al. 2020 Molecular and morphometric analyses identify new lineages within a large *Eumida* (Annelida) species complex. Zool. Scr. 49, 222–235. (doi: 10.1111/zsc.12397).

27. Blaxter M. 2016 Imagining Sisyphus happy: DNA barcoding and the unnamed majority. Philos. Trans. R. Soc. B Biol. Sci. 371, 20150329. (doi: 10.1098/rstb.2015.0329).

28. Costa F, Henzler C, Lunt D, Whiteley NM, Rock J. 2009 Probing marine *Gammarus* (Amphipoda) taxonomy with DNA barcodes. Syst. Biodivers. 7, 365–379. (doi: 10.1017/S1477200009990120).

29. Loeza-Quintana T et al. 2018 Recalibrating the molecular clock for Arctic marine invertebrates based on DNA barcodes. Genome 62, 1–17. (doi: 10.1139/gen-2018-0107).

30. Leese F, Kop A, Wägele JW, Held C. 2008 Cryptic speciation in a benthic isopod from Patagonian and Falkland Island waters and the impact of glaciations on its population structure. Front. Zool. 5, 19. (doi: 10.1186/1742-9994-5-19).

31. Markow TA, Pfeiler E. 2010 Mitochondrial DNA evidence for deep genetic divergences in allopatric populations of the rocky intertidal isopod *Ligia occidentalis* from the eastern Pacific. Mol. Phylogenet. Evol. 56, 468–473. (doi: 10.1016/j.ympev.2009.12.002).

32. Xavier R et al. 2011 Phylogeography of the marine isopod *Stenosoma nadejda* (Rezig, 1989) in North African Atlantic and western Mediterranean coasts reveals complex differentiation patterns and a new species. Biol. J. Linn. Soc. 104, 419–431. (doi: 10.1111/j.1095-8312.2011.01718.x).

33. Arnaud-Haond S et al. 2007 Vicariance patterns in the Mediterranean Sea: east–west cleavage and low dispersal in the endemic seagrass Posidonia oceanica. J. Biogeogr. 34, 963–976. (doi: 10.1111/j.1365-2699.2006.01671.x).

34. Varela AI, Haye PA. 2012 The marine brooder *Excirolana braziliensis* (Crustacea: Isopoda) is also a complex of cryptic species on the coast of Chile. Rev. Chil. Hist. Nat. 85, 495–502. (doi: 10.4067/S0716-078X2012000400011).

35. Sá-Pinto A, Branco M, Sayanda D, Alexandrino P. 2008 Patterns of colonization, evolution and gene flow in species of the genus *Patella* in the Macaronesian Islands. Mol. Ecol. 17, 519–532. (doi: 10.1111/j.1365-294X.2007.03563.x).

36. Xavier JR, Soest RWM. van Breeuwer AJ, Martin, AMF, Menken SBJ. 2010 Phylogeography, genetic diversity and structure of the poecilosclerid sponge *Phorbas fictitius* at oceanic islands. Contrib. Zool. 79, 119–129.

37. Quinteiro J, Rodríguez-Castro J, Rey-Méndez M, González-Henríquez N. 2020 Phylogeography of the insular populations of common octopus, *Octopus vulgaris* Cuvier, 1797, in the Atlantic Macaronesia. PLOS ONE 15, e0230294. (doi: 10.1371/journal.pone.0230294).

38. Domingues VS et al. 2008 Tropical fishes in a temperate sea: evolution of the wrasse *Thalassoma pavo* and the parrotfish *Sparisoma cretense* in the Mediterranean and the adjacent Macaronesian and Cape Verde Archipelagos. Mar. Biol. 154, 465–474. (doi: 10.1007/s00227-008-0941-z).

39. Beermann J et al. 2020 Ancient globetrotters—connectivity and putative native ranges of two cosmopolitan biofouling amphipods. PeerJ. 8, e9613. (doi: 10.7717/peerj.9613).

40. Thiel M, Gutow L. 2005 The ecology of rafting in the marine environment. II. The rafting organisms and community. 43, 279–418.

41. Waters JM, Fraser CI, Hewitt GM. 2013 Founder takes all: density-dependent processes structure biodiversity. Trends Ecol. Evol. 28, 78–85. (doi: 10.1016/j.tree.2012.08.024).

42. Torre G, Fernández-Lugo S, Guarino R, Fernández-Palacios JM. 2019 Network analysis by simulated annealing of taxa and islands of Macaronesia (North Atlantic Ocean). Ecography 42, 768–779. (doi: 0.1111/ecog.03909).

43. Hachich NF et al. 2015 Island biogeography: patterns of marine shallow-water organisms in the Atlantic Ocean. J. Biogeogr. 42, 1871–1882. (doi: 10.1111/jbi.12560).

44. Norder SJ et al. 2019 Beyond the Last Glacial Maximum: Island endemism is best explained by long-lasting archipelago configurations. Glob. Ecol. Biogeogr. 28, 184–197. (doi: 10.1111/geb.12835).

45. Jenkins TL, Castilho R, Stevens JR. 2018 Meta-analysis of northeast Atlantic marine taxa shows contrasting phylogeographic patterns following post-LGM expansions. PeerJ 6, e5684. (doi: 10.7717/peerj.5684).

46. Santos RS, Hawkins S, Monteiro LR, Alves M, Isidro EJ. 1995 Marine research, resources and conservation in the Azores. Aquat. Conserv. Mar. Freshw. Ecosyst. 5, 311–354.

47. Arístegui J et al. 2009 Sub-regional ecosystem variability in the Canary Current upwelling. Prog. Oceanogr. 83, 33–48. (doi: 10.1016/j.pocean.2009.07.031).

48. Hawkins SJ, Corte-Real HBSM, Pannacciulli FG, Weber LC, Bishop JDD. 2000 Thoughts on the ecology and evolution of the intertidal biota of the Azores and other Atlantic islands. Hydrobiologia 440, 3–17. (doi: 10.1023/A:1004118220083).

49. Xavier R, Branco M, dos Santos AM. 2016 Using a phylogeographic approach to investigate the diversity and determine the distributional range of an isopod (Crustacea: Peracarida), *Stenosoma nadejda* (Rezig, 1989) in the Atlantic-Mediterranean region. Hydrobiologia 768, 315–328. (doi: 10.1007/s10750-015-2559-8).

50. Cabezas MP, Cabezas P, Machordom A, Guerra-García JM. 2013 Hidden diversity and cryptic speciation refute cosmopolitan distribution in Caprella penantis (Crustacea: Amphipoda: Caprellidae). J. Zool. Syst. Evol. Res. 51, 85–99. (doi: 10.1111/jzs.12010).

51. Fernández-Palacios JM et al. 2011 A reconstruction of Palaeo-Macaronesia, with particular reference to the long-term biogeography of the Atlantic Island laurel forests. J. Biogeogr. 38, 226–246. (doi: 10.1111/j.1365-2699.2010.02427.x).

52. Radulovici AE, Archambault P, Dufresne F. 2010 DNA Barcodes for Marine Biodiversity: Moving Fast Forward? Diversity 2, 450–472. (doi: 10.3390/d2040450).

53. Hey J. 2006 On the failure of modern species concepts. Trends Ecol. Evol. 21, 447–450.

54. Coyne JA, Orr HA. 2004 Speciation. Oxford University Press.

55. Casacci LP, Barbero F, Balletto E. 2014 The “Evolutionarily Significant Unit” concept and its applicability in biological conservation. Ital. J. Zool. 81, 182–193. (doi: 10.1080/11250003.2013.870240).

56. Pante E et al. 2015 Species are hypotheses: avoid connectivity assessments based on pillars of sand. Mol. Ecol. 24, 525–544. (doi: 10.1111/mec.13048).

57. Allendorf F, Luikart G, Aitken S. 2012 Conservation and the Genetics of Populations. Wiley.

58. Frankham R. 1998 Inbreeding and Extinction: Island Populations. Conserv. Biol. 12, 665–675.

59. Korpinen S et al. 2021 Combined effects of human pressures on Europe’s marine ecosystems. Ambio. (doi:10.1007/s13280-020-01482-x).

60. Cacabelos E et al. 2020 Limited effects of marine protected areas on the distribution of invasive species, despite positive effects on diversity in shallow-water marine communities. Biol. Invasions 22, 1169–1179. (doi: 10.1007/s10530-019-02171-x).

61. Rogers AD et al. 2020 Critical Habitats and Biodiversity: Inventory, Thresholds and Governance. https://www.oceanpanel.org/blue-papers/critical-habitats-and-biodiversity-inventory-thresholds-and-governance.

62. Quinteiro J et al. 2012 Rede BANGEMAC: Banco genético marinho de Macaronésia (Memória técnica).

63. Vieira et al. 2021. Gaps in DNA sequence libraries for Macaronesian marine macroinvertebrates imply decades till completion and robust monitoring. Divers. Distrib. Accepted.

64. Hupało K et al. 2019 Persistence of phylogeographic footprints helps to understand cryptic diversity detected in two marine amphipods widespread in the Mediterranean basin. Mol. Phylogenet. Evol. 132, 53–66. (doi: 10.1016/j.ympev.2018.11.013).

65. Cheng J, Karambelkar B, Xie Y. 2018 leaflet: Create Interactive Web Maps with the JavaScript ‘Leaflet’ Library. R package version 2.0.1. https://CRAN.R-project.org/package=leaflet.

