## supplementary material for "Molecular evidence for sweeping discontinuity between peracarid (Crustacea) fauna of Macaronesian islands and nearby continental coasts: over fifty candidate endemic species"

### Material and Methods

#### *Specimen sampling and morphology based taxonomic identification*

Specimens were collected between 2011 and 2015 and sampled during low tide from marine intertidal rocky shores by scraping the algal cover or hand picking during low tide along the Northeast Atlantic coasts. After collection, specimens were preserved in 96% ethanol. The sampling locations are displayed in figure 1, and an interactive map with the sampling locations for each species is available at http://rpubs.com/Vieira/PeracaridaNE. The interactive map was created with the package 'leaflet' [1], through the software R 3.5.0 [2].

To ensure the correct taxonomic identification of the specimens, five steps were applied. Three based on morphology and two based on DNA barcoding (see below the subsection "Taxonomic identification with molecular tools"). The species' nomenclature used in this work complies with the accepted nomenclature used in World Register of Marine Species (WoRMS) and Integrated Taxonomic Information System (ITIS).

a) The first screening of the morphology-based taxonomic identification was supported in general keys for peracarids [3-9] and in the software package DELTA (DEscription Language for TAxonomy) with the interactive identification keys (INTKEY) for amphipods, isopods and tanaids [10-14];
b) second, updated and detailed identification keys for the genera *Gnathia* [15], *Dynamene* [16,17], *Campecopea* [8,18,19], *Cymodoce* [8,20,21], *Anthura* [22], *Apseudopis* [23,24], *Tanais* [25,26], *Caprella* [27-30], *Jassa* [31,32], *Elasmopus* [33-36], *Stenothoe* [37,38] and *Ampithoe* [39,40] were also used to accurately confirm the specimens identifications;
c) then, species checklists for the Northeast Atlantic and Macaronesia were used to verify/confirm the species presence and distribution in the locations where they were sampled [41-53].

#### *Genetic analysis and data treatment*

DNA extraction was carried out using the E.Z.N.A Mollusc DNA Kit (Omega Biotek), following the manufacturer's instructions. Depending on the specimen size, only a small amount of tissue or the whole animal was used. Then, the barcode region of the mitochondrial DNA gene cytochrome c oxidase subunit I (COI) was amplified in a MyCycler™ (Bio-Rad) thermal cycler using one of the three primer pairs: LCO1490/HCO2198 [54], LoboF1/LoboR1 [55] or LoboF1/ArR5 [56]. PCR thermal cycling conditions, primer sequences and references for each primer pair are available in Table S2. Each reaction contained 2.5 µl of 10× PCR buffer, 3 µl of 25 mM $MgCl_2$, 1 µl of 10 mM dNTP mixture, 0.2 µl of 5 U/µl of DNA Taq polymerase (ThermoScientific), 10 µM of each primer (1.25 µl for LoboF1/LoboR1; 0.5 µl for LCO1490/HCO2198; 0.55 µl for ArR5), 2-4 µl of DNA template and completed with sterile milli Q-grade water to make up a total volume of 25 µl.

Amplification success was verified in a 1.5% agarose gel. DNA templates were then purified (with Roche purification kit according to manufacturer instructions) and sequenced bidirectionally in an external sequencing service supplier (Macrogen Europe, Netherlands).

Each trace file was edited individually and manually, unreadable zones and primers removed, and ambiguous bases corrected. The resultant sequences were aligned using Clustal W [57] implemented in MEGA 7.0 [58] and inspected for eventual anomalies, such as stop codons or indels in DNASP 5.10 [59]. Sequences of different length were obtained depending on the primer used (Table S2). In the end, a common fragment of 520 base pair (bp) obtained from all sequences was used.

*Taxonomic identification with molecular tools*

After obtaining the COI sequences for each specimen, two additional steps were applied based on the molecular data to confirm and support the morphological identifications:

d) GenBank BLASTn search [60] and BOLD Identification System tool (BOLD-IDS) [61] were used to search for similarity to confirm the target taxa;
e) and finally, sister taxa (see Table S3 for the list of species, number of specimens and source) commonly found in Madeira, Selvagens, Azores, Canaries, Iberian Peninsula or Morocco [41–53], were amplified by us (as above) or mined from BOLD/GenBank and used to construct phylogenetic trees to verify the clustering of the different taxa. The Bayesian inference (BI) was conducted in MrBayes 3.2 [62] to build the Bayesian tree for each order separately. The BI topologies were constructed choosing GTR+G+I as best-fitting model of nucleotide substitution based on its Bayesian Information Criterion as implemented in MEGA 7.0 [58]. Runs were conducted with 7 x $10^6$ generations each. Parameters were sampled every 1 x $10^2$ generations and then a burn-in of 25% was applied.

*Molecular species delimitation*

Five methods were used to determine the number of MOTUs. First, we used two distance-based barcode gap approaches. The Automatic Barcode Gap Discovery (ABGD) species delineation tool was performed on a web interface (http://wwwabi.snv.jussieu.fr/public/abgd/abgdweb.html) with default settings for the Kimura-2-parameter (K2P) distance matrix. This tool, based on the barcode gap detection (i.e., break between the distribution of intraspecific and interspecific distances of the barcode region), sorts the sequences into hypothetical species [63]. Then, the Cluster Sequences tool implemented in BOLD v4 (http://v4.boldsystems.org) [61], was applied. This approach clusters barcode sequences algorithmically to calculate MOTUs that show high concordance to species [64].

Then, two tree-based methods were applied: bPTP [65] that requires a non-ultrametric tree and GMYC single-threshold model [66] that requires an ultrametric tree. The bPTP method incorporates the number of substitutions in the model of speciation and assumes that the probability that a substitution gives rise to a speciation event follows a Poisson distribution. The branch lengths of the input tree are supposed to be generated by two independent Poisson process classes, one corresponding to speciation and the other to coalescence [65]. The GMYC method is based on the examination of the branching patterns in an ultrametric tree, and the recognition of the transitions from branching patterns attributable to speciation (one lineage per species) to those that can be attributed to intra-species coalescent process (multiple lineages per species). We only applied the single threshold variant of this method [67] because it outperforms the multi-threshold version [68], as suggested by Fujisawa & Barraclough [66]. To apply the bPTP method, the web server bPTP was used (http://species.h-its.org/ptp) [65]. Maximum-likelihood (ML) trees for COI (1 x $10^3$ bootstrap) was constructed with MEGA 7.0 [58] and used as input. Evolutionary models were selected using also MEGA 7.0 [58] under the corrected Akaike information criterion. Species delimitations were performed using 5 x $10^5$ Markov chain Monte Carlo iterations (MCMC) with a 25% burn-in. Since the GMYC method requires an ultrametric tree, we first calculated a Bayesian ultrametric phylogenetic tree for each order. The tree was generated in BEAST 2.4.6 [69] with the appropriate best model (calculated as above, but based on Bayesian Information Criterion), and four independent runs for 7 x $10^7$ MCMC generations, sampled every 1 x $10^4$ generations, were performed. Convergence of the parameters was evaluated (accordingly with BEAST recommendation) using Tracer 1.6 software [70]. The consensus tree was annotated using TreeAnnotator 2.4.6 [69]. The consensus tree was loaded into the package 'SPLITS' (Species Limits by Threshold Statistics) [71] in R 3.5.0 [2] and analysed using the single-threshold model.

Finally, the 95% statistical parsimony connection limit was applied with TCS 1.21 [72]. This is a common method derived from population genetics to visualize possible intraspecific relationships. Sequences are assigned to networks connected by changes, which are non-homoplastic with a certain probability. Even though this is not equivalent to defining species boundaries, statistical parsimony has also been applied successfully to delimit candidate species before (e.g. [67,73]).

*Network analysis*

The tests with locations (i.e. islands plus Iberian Peninsula and Morocco) and regions (MACA, IP, AZ, MORO) retrieved the networks as small (i.e., number of nodes lower than 500) with the mixing parameter μ (reflects the accuracy of the community detection algorithms) lower than 0.5. For small networks with μ lower than 0.5 (see Yang *et al.* [74] for details), the following algorithms can be accepted: Infomap, Label propagation, Multilevel, Walktrap, Spinglass and the Edge betweenness.

**Table S1.** List of peracaridean species used in the molecular species delineation, with respective source, sampling location and number of individuals used.

| Order | Species | Source | Country/Island | Site | Region | Latitude | Longitude | N |
|---|---|---|---|---|---|---|---|---|
| **AMPHIPODA** | *Ampithoe helleri* | 55,75 | Portugal | Viana do Castelo | IP | 41.689194 | -8.84787 | 8 |
| **AMPHIPODA** | *Ampithoe helleri* | This study | Gran Canaria | Bañaderos | MACA | 28.149658 | -15.54018 | 3 |
| **AMPHIPODA** | *Ampithoe helleri* | This study | Spain | Barizo | IP | 43.322113 | -8.872784 | 2 |
| **AMPHIPODA** | *Ampithoe helleri* | This study | La Palma | El Faro | MACA | 28.457545 | -17.85034 | 2 |
| **AMPHIPODA** | *Ampithoe ramondi* | This study | Tenerife | Mal Paso | MACA | 28.416147 | -16.298656 | 5 |
| **AMPHIPODA** | *Ampithoe ramondi* | This study | Portugal | Ingrina | IP | 37.045257 | -8.878047 | 1 |
| **AMPHIPODA** | *Ampithoe ramondi* | This study | Portugal | Arrifes | IP | 37.076052 | -8.27678 | 1 |
| **AMPHIPODA** | *Ampithoe ramondi* | This study | Portugal | Dona Ana | IP | 37.086969 | -8.667716 | 2 |
| **AMPHIPODA** | *Ampithoe ramondi* | This study | Gran Canaria | Bañaderos | MACA | 28.149658 | -15.54018 | 1 |
| **AMPHIPODA** | *Ampithoe ramondi* | This study | Madeira | Ponta da Cruz | MACA | 32.633123 | -16.943643 | 1 |
| **AMPHIPODA** | *Ampithoe ramondi* | This study | Santa Maria | Praia Formosa | AZ | 36.949917 | -25.094989 | 1 |
| **AMPHIPODA** | *Ampithoe ramondi* | This study | Santa Maria | São Lourenço | AZ | 36.988172 | -25.054211 | 1 |
| **AMPHIPODA** | *Ampithoe riedli* | This study | Portugal | Ingrina | IP | 37.045257 | -8.878047 | 3 |
| **AMPHIPODA** | *Ampithoe riedli* | This study | Morocco | Arzila | MOCO | 35.458006 | -6.047981 | 3 |
| **AMPHIPODA** | *Ampithoe riedli* | This study | Madeira | Ponta da Cruz | MACA | 32.633123 | -16.943643 | 2 |
| **AMPHIPODA** | *Ampithoe riedli* | This study | La Palma | La Fajana | MACA | 28.842276 | -17.794324 | 1 |
| **AMPHIPODA** | *Apohyale perieri* | 76 | Gran Canaria | Agaete | MACA | 28.163186 | -15.699269 | 1 |
| **AMPHIPODA** | *Apohyale perieri* | 76 | La Palma | El Faro | MACA | 28.457545 | -17.85034 | 3 |
| **AMPHIPODA** | *Apohyale perieri* | 76 | La Palma | La Fajana | MACA | 28.842276 | -17.794324 | 1 |
| **AMPHIPODA** | *Apohyale perieri* | 76 | Madeira | Ponta da Cruz | MACA | 32.633123 | -16.943643 | 3 |
| **AMPHIPODA** | *Apohyale perieri* | 76 | Portugal | Arrifes | IP | 37.076052 | -8.27678 | 1 |
| **AMPHIPODA** | *Apohyale perieri* | 76 | Portugal | Buarcos | IP | 40.175976 | -8.900572 | 2 |
| **AMPHIPODA** | *Apohyale perieri* | 76 | Portugal | Agudela | IP | 41.240725 | -8.727547 | 1 |
| **AMPHIPODA** | *Apohyale perieri* | 76 | Portugal | São Pedro Moel | IP | 39.758015 | -9.033165 | 1 |
| **AMPHIPODA** | *Apohyale perieri* | 76 | São Miguel | Ponta da Ferreirinha | AZ | 37.861000 | -25.8548 | 1 |
| **AMPHIPODA** | *Apohyale perieri* | 76 | Spain | Muxía | IP | 43.092831 | -9.223431 | 1 |
| **AMPHIPODA** | *Apohyale perieri* | 76 | Spain | Barizo | IP | 43.322113 | -8.872784 | 3 |
| **AMPHIPODA** | *Apohyale perieri* | 76 | Spain | Pedreira | IP | 43.55617 | -8.274942 | 3 |
| **AMPHIPODA** | *Apohyale stebbingi* | 76 | Gran Canaria | Agaete | MACA | 28.163186 | -15.699269 | 2 |
| **AMPHIPODA** | *Apohyale stebbingi* | 76 | Gran Canaria | Playa Melenara | MACA | 27.988891 | -15.370485 | 1 |
| **AMPHIPODA** | *Apohyale stebbingi* | 76 | La Palma | El Faro | MACA | 28.457545 | -17.85034 | 4 |
| **AMPHIPODA** | *Apohyale stebbingi* | 76 | La Palma | La Salemera | MACA | 28.577985 | -17.760556 | 2 |
| **AMPHIPODA** | *Apohyale stebbingi* | 76 | Madeira | Ponta da Cruz | MACA | 32.633123 | -16.943643 | 4 |
| **AMPHIPODA** | *Apohyale stebbingi* | 76 | Madeira | Reis Magos | MACA | 32.646111 | -16.824167 | 1 |
| **AMPHIPODA** | *Apohyale stebbingi* | 76 | Morocco | Arzila | MOCO | 35.458006 | -6.047981 | 4 |

| Order | Species | Source | Country/Island | Site | Region | Latitude | Longitude | N |
|---|---|---|---|---|---|---|---|---|
| **AMPHIPODA** | *Apohyale stebbingi* | 76 | Portugal | Ingrina | IP | 37.045257 | -8.878047 | 1 |
| **AMPHIPODA** | *Apohyale stebbingi* | 76 | Portugal | Arrifes | IP | 37.076052 | -8.276780 | 2 |
| **AMPHIPODA** | *Apohyale stebbingi* | 76 | Portugal | Dona Ana | IP | 37.086969 | -8.667716 | 1 |
| **AMPHIPODA** | *Apohyale stebbingi* | 76 | Portugal | Agudela | IP | 41.240725 | -8.727547 | 1 |
| **AMPHIPODA** | *Apohyale stebbingi* | 76 | Portugal | Peniche | IP | 39.372433 | -9.377551 | 2 |
| **AMPHIPODA** | *Apohyale stebbingi* | 76 | Portugal | São Pedro Moel | IP | 39.758015 | -9.033165 | 2 |
| **AMPHIPODA** | *Apohyale stebbingi* | 76 | Santa Maria | Praia Formosa | AZ | 36.949917 | -25.094989 | 1 |
| **AMPHIPODA** | *Apohyale stebbingi* | 76 | São Miguel | Mosteiros | AZ | 37.900153 | -25.817875 | 1 |
| **AMPHIPODA** | *Apohyale stebbingi* | 76 | São Miguel | Ponta da Ferreirinha | AZ | 37.861000 | -25.854800 | 3 |
| **AMPHIPODA** | *Apohyale stebbingi* | 76 | Spain | Muxía | IP | 43.092831 | -9.223431 | 1 |
| **AMPHIPODA** | *Apohyale stebbingi* | 76 | Spain | Pedreira | IP | 43.556170 | -8.274942 | 2 |
| **AMPHIPODA** | *Apohyale stebbingi* | 76 | Tenerife | Los Cristianos | MACA | 28.044714 | -16.711856 | 1 |
| **AMPHIPODA** | *Caprella acanthifera* | This Study | El Hierro | Arenas Blancas | MACA | 27.767189 | -18.121308 | 1 |
| **AMPHIPODA** | *Caprella acanthifera* | This Study | Gran Canaria | Agaete | MACA | 28.163186 | -15.699269 | 1 |
| **AMPHIPODA** | *Caprella acanthifera* | This Study | La Palma | El Faro | MACA | 28.457545 | -17.850340 | 2 |
| **AMPHIPODA** | *Caprella acanthifera* | This Study | La Palma | La Salemera | MACA | 28.577985 | -17.760556 | 1 |
| **AMPHIPODA** | *Caprella acanthifera* | This Study | Madeira | Ponta da Cruz | MACA | 32.633123 | -16.943643 | 2 |
| **AMPHIPODA** | *Caprella acanthifera* | This Study | Madeira | Reis Magos | MACA | 32.646111 | -16.824167 | 1 |
| **AMPHIPODA** | *Caprella acanthifera* | This Study | Morocco | El Jadida | MOCO | 33.264036 | -8.510717 | 2 |
| **AMPHIPODA** | *Caprella acanthifera* | 75 | Portugal | Viana do Castelo | IP | 41.689194 | -8.847870 | 3 |
| **AMPHIPODA** | *Caprella acanthifera* | This Study | Portugal | Buarcos | IP | 40.175976 | -8.900572 | 2 |
| **AMPHIPODA** | *Caprella acanthifera* | This Study | São Miguel | Ribeira Chã | AZ | 37.715417 | -25.486836 | 3 |
| **AMPHIPODA** | *Caprella acanthifera* | This Study | Tenerife | Los Cristianos | MACA | 28.044714 | -16.711856 | 1 |
| **AMPHIPODA** | *Elasmopus pectenicrus* | This study | Tenerife | Mal Paso | MACA | 28.416147 | -16.298656 | 2 |
| **AMPHIPODA** | *Elasmopus pectenicrus* | This study | Porto Santo | Porto dos Frades | MACA | 33.072575 | -16.295666 | 2 |
| **AMPHIPODA** | *Elasmopus pectenicrus* | This study | Portugal | Arrifes | IP | 37.076052 | -8.276780 | 1 |
| **AMPHIPODA** | *Elasmopus pectenicrus* | This study | Portugal | Dona Ana | IP | 37.086969 | -8.667716 | 2 |
| **AMPHIPODA** | *Elasmopus pectenicrus* | This study | Morocco | Akhfenir | MOCO | 28.097524 | -12.050701 | 3 |
| **AMPHIPODA** | *Elasmopus pectenicrus* | This study | Madeira | Ponta da Cruz | MACA | 32.633123 | -16.943643 | 3 |
| **AMPHIPODA** | *Elasmopus pectenicrus* | This study | Madeira | Reis Magos | MACA | 32.646111 | -16.824167 | 1 |
| **AMPHIPODA** | *Jassa herdmani* | This study | Madeira | Ponta da Cruz | MACA | 32.633123 | -16.943643 | 2 |
| **AMPHIPODA** | *Jassa herdmani* | This study | Porto Santo | Porto dos Frades | MACA | 33.072575 | -16.295666 | 1 |
| **AMPHIPODA** | *Jassa herdmani* | 55,75 | Portugal | Viana do Castelo | IP | 41.689194 | -8.847870 | 2 |
| **AMPHIPODA** | *Jassa herdmani* | This study | Portugal | Buarcos | IP | 40.175976 | -8.900572 | 3 |
| **AMPHIPODA** | *Jassa herdmani* | This study | São Miguel | Ribeira Chã | AZ | 37.715417 | -25.486836 | 3 |
| **AMPHIPODA** | *Podocerus variegatus* | This study | La Palma | El Faro | MACA | 28.457545 | -17.850340 | 1 |
| **AMPHIPODA** | *Podocerus variegatus* | This study | La Palma | La Fajana | MACA | 28.842276 | -17.794324 | 2 |
| **AMPHIPODA** | *Podocerus variegatus* | This study | Porto Santo | Porto dos Frades | MACA | 33.072575 | -16.295666 | 1 |

| Order | Species | Source | Country/Island | Site | Region | Latitude | Longitude | N |
|---|---|---|---|---|---|---|---|---|
| **AMPHIPODA** | *Podocerus variegatus* | This study | Spain | Muxía | IP | 43.092831 | -9.223431 | 2 |
| **AMPHIPODA** | *Podocerus variegatus* | This study | Spain | Barizo | IP | 43.322113 | -8.872784 | 2 |
| **AMPHIPODA** | *Podocerus variegatus* | This study | Spain | Pedreira | IP | 43.556170 | -8.274942 | 1 |
| **AMPHIPODA** | *Protohyale schmidtii* | 76 | Gran Canaria | Bañaderos | MACA | 28.149658 | -15.540180 | 3 |
| **AMPHIPODA** | *Protohyale schmidtii* | 76 | La Palma | El Faro | MACA | 28.457545 | -17.850340 | 1 |
| **AMPHIPODA** | *Protohyale schmidtii* | 76 | La Palma | La Salemera | MACA | 28.577985 | -17.760556 | 1 |
| **AMPHIPODA** | *Protohyale schmidtii* | 76 | La Palma | La Fajana | MACA | 28.842276 | -17.794324 | 1 |
| **AMPHIPODA** | *Protohyale schmidtii* | 76 | Madeira | Ponta da Cruz | MACA | 32.633123 | -16.943643 | 1 |
| **AMPHIPODA** | *Protohyale schmidtii* | 76 | Morocco | Akhfenir | MOCO | 28.097524 | -12.050701 | 2 |
| **AMPHIPODA** | *Protohyale schmidtii* | 76 | Morocco | Tarfaya | MOCO | 27.917817 | -12.961147 | 1 |
| **AMPHIPODA** | *Protohyale schmidtii* | 76 | Porto Santo | Porto dos Frades | MACA | 33.072575 | -16.295666 | 1 |
| **AMPHIPODA** | *Protohyale schmidtii* | 76 | Portugal | Arrifes | IP | 37.076052 | -8.276780 | 4 |
| **AMPHIPODA** | *Protohyale schmidtii* | 76 | Portugal | Buarcos | IP | 40.175976 | -8.900572 | 1 |
| **AMPHIPODA** | *Protohyale schmidtii* | 76 | Portugal | Peniche | IP | 39.372433 | -9.377551 | 1 |
| **AMPHIPODA** | *Protohyale schmidtii* | 76 | Santa Maria | Praia Formosa | AZ | 36.949917 | -25.094989 | 2 |
| **AMPHIPODA** | *Protohyale schmidtii* | 76 | Santa Maria | São Lourenço | AZ | 36.988172 | -25.054211 | 2 |
| **AMPHIPODA** | *Protohyale schmidtii* | 76 | São Miguel | Ribeira Chã | AZ | 37.715417 | -25.486836 | 1 |
| **AMPHIPODA** | *Protohyale schmidtii* | 76 | São Miguel | Mosteiros | AZ | 37.900153 | -25.817875 | 1 |
| **AMPHIPODA** | *Protohyale schmidtii* | 76 | El Hierro | Los Sargos | MACA | 27.784739 | -18.011569 | 1 |
| **AMPHIPODA** | *Protohyale schmidtii* | 76 | Spain | Muxía | IP | 43.092831 | -9.223431 | 2 |
| **AMPHIPODA** | *Protohyale schmidtii* | 76 | Spain | Barizo | IP | 43.322113 | -8.872784 | 3 |
| **AMPHIPODA** | *Protohyale schmidtii* | 76 | Spain | Pedreira | IP | 43.556170 | -8.274942 | 2 |
| **AMPHIPODA** | *Protohyale schmidtii* | 76 | Tenerife | Mal Paso | MACA | 28.416147 | -16.298656 | 2 |
| **AMPHIPODA** | *Protohyale schmidtii* | 76 | Tenerife | Los Cristianos | MACA | 28.044714 | -16.711856 | 1 |
| **AMPHIPODA** | *Quadrimaera inaequipes* | 55,75 | Portugal | Viana do Castelo | IP | 41.689194 | -8.847870 | 8 |
| **AMPHIPODA** | *Quadrimaera inaequipes* | This study | Gran Canaria | Bañaderos | MACA | 28.149658 | -15.540180 | 1 |
| **AMPHIPODA** | *Quadrimaera inaequipes* | This study | La Palma | El Faro | MACA | 28.457545 | -17.850340 | 1 |
| **AMPHIPODA** | *Quadrimaera inaequipes* | This study | La Palma | La Salemera | MACA | 28.577985 | -17.760556 | 1 |
| **AMPHIPODA** | *Quadrimaera inaequipes* | This study | Madeira | Ponta da Cruz | MACA | 32.633123 | -16.943643 | 3 |
| **AMPHIPODA** | *Quadrimaera inaequipes* | This study | La Palma | La Fajana | MACA | 28.842276 | -17.794324 | 1 |
| **AMPHIPODA** | *Serejohyale spinidactylus* | 76 | El Hierro | Los Sargos | MACA | 27.784739 | -18.011569 | 1 |
| **AMPHIPODA** | *Serejohyale spinidactylus* | 76 | Gran Canaria | Bañaderos | MACA | 28.149658 | -15.54018 | 3 |
| **AMPHIPODA** | *Serejohyale spinidactylus* | 76 | Gran Canaria | Agaete | MACA | 28.163186 | -15.699269 | 1 |
| **AMPHIPODA** | *Serejohyale spinidactylus* | 76 | Gran Canaria | Playa Melenara | MACA | 27.988891 | -15.370485 | 2 |
| **AMPHIPODA** | *Serejohyale spinidactylus* | 76 | La Palma | La Salemera | MACA | 28.577985 | -17.760556 | 4 |
| **AMPHIPODA** | *Serejohyale spinidactylus* | 76 | La Palma | La Fajana | MACA | 28.842276 | -17.794324 | 2 |
| **AMPHIPODA** | *Serejohyale spinidactylus* | 76 | Selvagens | Selvagem Grande | MACA | 30.141158 | -15.870064 | 1 |
| **AMPHIPODA** | *Serejohyale spinidactylus* | 76 | Madeira | Ponta da Cruz | MACA | 32.633123 | -16.943643 | 1 |

| Order | Species | Source | Country/Island | Site | Region | Latitude | Longitude | N |
|---|---|---|---|---|---|---|---|---|
| **AMPHIPODA** | *Serejohyale spinidactylus* | 76 | Madeira | Reis Magos | MACA | 32.646111 | -16.824167 | 2 |
| **AMPHIPODA** | *Serejohyale spinidactylus* | 76 | São Miguel | Mosteiros | AZ | 37.900153 | -25.817875 | 1 |
| **AMPHIPODA** | *Serejohyale spinidactylus* | 76 | São Miguel | Ponta da Ferreirinha | AZ | 37.861000 | -25.854800 | 2 |
| **AMPHIPODA** | *Serejohyale spinidactylus* | 76 | Spain | Muxía | IP | 43.092831 | -9.223431 | 3 |
| **AMPHIPODA** | *Serejohyale spinidactylus* | 76 | Spain | Barizo | IP | 43.322113 | -8.872784 | 2 |
| **AMPHIPODA** | *Stenothoe monoculoides* | 77 | North Sea | Helgoland | - | 54.171000 | 7.889000 | 4 |
| **AMPHIPODA** | *Stenothoe monoculoides* | This study | Tenerife | Mal Paso | MACA | 28.416147 | -16.298656 | 3 |
| **ISOPODA** | *Anthura gracilis* | This study | Tenerife | Los Cristianos | MACA | 28.044714 | -16.711856 | 1 |
| **ISOPODA** | *Anthura gracilis* | This study | Porto Santo | Porto dos Frades | MACA | 33.072575 | -16.295666 | 1 |
| **ISOPODA** | *Anthura gracilis* | This study | Selvagens | Selvagem Grande | MACA | 30.141158 | -15.870064 | 1 |
| **ISOPODA** | *Anthura gracilis* | This study | Gran Canaria | Agaete | MACA | 28.163186 | -15.699269 | 1 |
| **ISOPODA** | *Anthura gracilis* | This study | Spain | Barizo | IP | 43.322113 | -8.872784 | 1 |
| **ISOPODA** | *Anthura gracilis* | This study | Morocco | Arzila | MOCO | 35.458006 | -6.047981 | 2 |
| **ISOPODA** | *Anthura gracilis* | This study | Terceira | Porto Martins | AZ | 38.683328 | -27.057522 | 1 |
| **ISOPODA** | *Anthura gracilis* | This study | Portugal | Viana do Castelo | IP | 41.689194 | -8.84787 | 2 |
| **ISOPODA** | *Anthura gracilis* | This study | São Miguel | Ribeira Chã | AZ | 37.715417 | -25.486836 | 3 |
| **ISOPODA** | *Anthura gracilis* | This study | La Palma | La Fajana | MACA | 28.842276 | -17.794324 | 1 |
| **ISOPODA** | *Campecopea lusitanica* | This study | Porto Santo | Porto dos Frades | MACA | 33.072575 | -16.295666 | 2 |
| **ISOPODA** | *Campecopea lusitanica* | This study | Gran Canaria | Bañaderos | MACA | 28.149658 | -15.540180 | 1 |
| **ISOPODA** | *Campecopea lusitanica* | This study | Portugal | Peniche | IP | 39.372433 | -9.377551 | 1 |
| **ISOPODA** | *Campecopea lusitanica* | This study | La Palma | El Faro | MACA | 28.457545 | -17.85034 | 1 |
| **ISOPODA** | *Campecopea lusitanica* | This study | Spain | Pedreira | IP | 43.556170 | -8.274942 | 3 |
| **ISOPODA** | *Campecopea lusitanica* | This study | La Palma | La Fajana | MACA | 28.842276 | -17.794324 | 1 |
| **ISOPODA** | *Cymodoce truncata* | This study | Porto Santo | Porto dos Frades | MACA | 33.072575 | -16.295666 | 2 |
| **ISOPODA** | *Cymodoce truncata* | This study | Spain | Muxía | IP | 43.092831 | -9.223431 | 1 |
| **ISOPODA** | *Cymodoce truncata* | This study | Portugal | Vale dos Homens | IP | 37.37140 | -8.834500 | 1 |
| **ISOPODA** | *Cymodoce truncata* | This study | Portugal | Peniche | IP | 39.372433 | -9.377551 | 3 |
| **ISOPODA** | *Cymodoce truncata* | This study | Madeira | Ponta da Cruz | MACA | 32.633123 | -16.943643 | 1 |
| **ISOPODA** | *Cymodoce truncata* | This study | Terceira | Porto Martins | AZ | 38.683328 | -27.057522 | 1 |
| **ISOPODA** | *Cymodoce truncata* | This study | La Palma | La Fajana | MACA | 28.842276 | -17.794324 | 2 |
| **ISOPODA** | *Dynamene edwardsi* | 78 | El Hierro | Arenas Blancas | MACA | 27.767189 | -18.121308 | 3 |
| **ISOPODA** | *Dynamene edwardsi* | 78 | El Hierro | Los Sargos | MACA | 27.784739 | -18.011569 | 3 |
| **ISOPODA** | *Dynamene edwardsi* | 78 | Gran Canaria | Bañaderos | MACA | 28.149658 | -15.540180 | 6 |
| **ISOPODA** | *Dynamene edwardsi* | 78 | Gran Canaria | Agaete | MACA | 28.163186 | -15.699269 | 6 |
| **ISOPODA** | *Dynamene edwardsi* | 78 | Gran Canaria | Playa Melenara | MACA | 27.988891 | -15.370485 | 5 |
| **ISOPODA** | *Dynamene edwardsi* | 78 | La Palma | El Faro | MACA | 28.457545 | -17.850340 | 6 |
| **ISOPODA** | *Dynamene edwardsi* | 78 | La Palma | La Salemera | MACA | 28.577985 | -17.760556 | 5 |
| **ISOPODA** | *Dynamene edwardsi* | 78 | La Palma | La Fajana | MACA | 28.842276 | -17.794324 | 5 |

| Order | Species | Source | Country/Island | Site | Region | Latitude | Longitude | N |
|---|---|---|---|---|---|---|---|---|
| **ISOPODA** | *Dynamene edwardsi* | 78 | Madeira | Ponta da Cruz | MACA | 32.633123 | -16.943643 | 5 |
| **ISOPODA** | *Dynamene edwardsi* | 78 | Madeira | Reis Magos | MACA | 32.646111 | -16.824167 | 5 |
| **ISOPODA** | *Dynamene edwardsi* | 78 | Morocco | El Jadida | MOCO | 33.264036 | -8.510717 | 1 |
| **ISOPODA** | *Dynamene edwardsi* | 78 | Morocco | Arzila | MOCO | 35.458006 | -6.047981 | 4 |
| **ISOPODA** | *Dynamene edwardsi* | 78 | Morocco | Tarfaya | MOCO | 27.917817 | -12.961147 | 4 |
| **ISOPODA** | *Dynamene edwardsi* | 78 | Porto Santo | Porto dos Frades | MACA | 33.072575 | -16.295666 | 5 |
| **ISOPODA** | *Dynamene edwardsi* | 78 | Portugal | Ingrina | IP | 37.045257 | -8.878047 | 5 |
| **ISOPODA** | *Dynamene edwardsi* | 78 | Portugal | Arrifes | IP | 37.076052 | -8.276780 | 4 |
| **ISOPODA** | *Dynamene edwardsi* | 78 | Portugal | Dona Ana | IP | 37.086969 | -8.667716 | 5 |
| **ISOPODA** | *Dynamene edwardsi* | 78 | Portugal | Peniche | IP | 39.372433 | -9.377551 | 4 |
| **ISOPODA** | *Dynamene edwardsi* | 78 | Portugal | Sines | IP | 37.960884 | -8.887296 | 1 |
| **ISOPODA** | *Dynamene edwardsi* | 78 | São Miguel | Mosteiros | AZ | 37.900153 | -25.817875 | 1 |
| **ISOPODA** | *Dynamene edwardsi* | 78 | Selvagens | Selvagem Pequena | MACA | 30.033233 | -16.016675 | 2 |
| **ISOPODA** | *Dynamene edwardsi* | 78 | Selvagens | Selvagem Grande | MACA | 30.141158 | -15.870064 | 4 |
| **ISOPODA** | *Dynamene edwardsi* | 78 | Spain | Muxía | IP | 43.092831 | -9.223431 | 1 |
| **ISOPODA** | *Dynamene edwardsi* | 78 | Tenerife | Los Cristianos | MACA | 28.044714 | -16.711856 | 4 |
| **ISOPODA** | *Dynamene edwardsi* | 78 | Tenerife | Mal Paso | MACA | 28.416147 | -16.298656 | 5 |
| **ISOPODA** | *Gnathia maxillaris* | This study | Gran Canaria | Agaete | MACA | 28.163186 | -15.699269 | 2 |
| **ISOPODA** | *Gnathia maxillaris* | This study | La Palma | El Faro | MACA | 28.457545 | -17.850340 | 1 |
| **ISOPODA** | *Gnathia maxillaris* | This study | La Palma | La Fajana | MACA | 28.842276 | -17.794324 | 2 |
| **ISOPODA** | *Gnathia maxillaris* | This study | Porto Santo | Porto dos Frades | MACA | 33.072575 | -16.295666 | 1 |
| **ISOPODA** | *Gnathia maxillaris* | This study | Portugal | Ingrina | IP | 37.045257 | -8.878047 | 1 |
| **ISOPODA** | *Gnathia maxillaris* | This study | Portugal | Buarcos | IP | 40.175976 | -8.900572 | 2 |
| **ISOPODA** | *Gnathia maxillaris* | This study | Spain | Pedreira | IP | 43.556170 | -8.274942 | 1 |
| **ISOPODA** | *Janira maculosa* | This study | Portugal | Dona Ana | IP | 37.086969 | -8.667716 | 2 |
| **ISOPODA** | *Janira maculosa* | This study | Spain | Muxía | IP | 43.092831 | -9.223431 | 1 |
| **ISOPODA** | *Janira maculosa* | This study | La Palma | La Salemera | MACA | 28.577985 | -17.760556 | 2 |
| **ISOPODA** | *Janira maculosa* | This study | La Palma | La Fajana | MACA | 28.842276 | -17.794324 | 1 |
| **ISOPODA** | *Joeropsis brevicornis* | This study | Tenerife | Los Cristianos | MACA | 28.044714 | -16.711856 | 2 |
| **ISOPODA** | *Joeropsis brevicornis* | This study | Portugal | Dona Ana | IP | 37.086969 | -8.667716 | 2 |
| **ISOPODA** | *Joeropsis brevicornis* | This study | Spain | Barizo | IP | 43.322113 | -8.872784 | 1 |
| **ISOPODA** | *Joeropsis brevicornis* | This study | La Palma | El Faro | MACA | 28.457545 | -17.85034 | 2 |
| **ISOPODA** | *Joeropsis brevicornis* | This study | Madeira | Reis Magos | MACA | 32.646111 | -16.824167 | 2 |
| **TANAIDACEA** | *Apseudopsis latreilii* | This study | Porto Santo | Porto dos Frades | MACA | 33.072575 | -16.295666 | 1 |
| **TANAIDACEA** | *Apseudopsis latreilii* | This study | Portugal | Dona Ana | IP | 37.086969 | -8.667716 | 3 |
| **TANAIDACEA** | *Apseudopsis latreilii* | This study | Gran Canaria | Agaete | MACA | 28.163186 | -15.699269 | 2 |
| **TANAIDACEA** | *Tanais dulongii* | This study | La Palma | El Faro | MACA | 28.457545 | -17.85034 | 1 |
| **TANAIDACEA** | *Tanais dulongii* | This study | La Palma | La Salemera | MACA | 28.577985 | -17.760556 | 2 |

| Order | Species | Source | Country/Island | Site | Region | Latitude | Longitude | N |
|---|---|---|---|---|---|---|---|---|
| **TANAIDACEA** | *Tanais dulongii* | This study | Madeira | Ponta da Cruz | MACA | 32.633123 | -16.943643 | 3 |
| **TANAIDACEA** | *Tanais dulongii* | This study | Morocco | El Jadida | MOCO | 33.264036 | -8.510717 | 1 |
| **TANAIDACEA** | *Tanais dulongii* | This study | Morocco | Arzila | MOCO | 35.458006 | -6.047981 | 1 |
| **TANAIDACEA** | *Tanais dulongii* | This study | Portugal | Viana do Castelo | IP | 41.689194 | -8.847870 | 2 |
| **TANAIDACEA** | *Tanais dulongii* | This study | Portugal | Ingrina | IP | 37.045257 | -8.878047 | 1 |
| **TANAIDACEA** | *Tanais dulongii* | This study | Portugal | Peniche | IP | 39.372433 | -9.377551 | 1 |
| **TANAIDACEA** | *Tanais dulongii* | This study | Portugal | Berlengas | IP | 39.411773 | -9.510989 | 1 |
| **TANAIDACEA** | *Tanais dulongii* | This study | Spain | Barizo | IP | 43.322113 | -8.872784 | 1 |
| **TANAIDACEA** | *Tanais grimaldii* | This study | Selvagens | Selvagem Pequena | MACA | 30.033233 | -16.016675 | 1 |
| **TANAIDACEA** | *Tanais grimaldii* | This study | Porto Santo | Porto dos Frades | MACA | 33.072575 | -16.295666 | 2 |
| **TANAIDACEA** | *Tanais grimaldii* | This study | Selvagens | Selvagem Grande | MACA | 30.141158 | -15.870064 | 2 |
| **TANAIDACEA** | *Tanais grimaldii* | This study | Spain | Barizo | IP | 43.322113 | -8.872784 | 2 |
| **TANAIDACEA** | *Tanais grimaldii* | This study | São Miguel | Ribeira Chã | AZ | 37.715417 | -25.486836 | 2 |

Az- Azores; MACA - Webbnesia; MORO – Morocco, IP - Iberian Peninsula.

**Table S2.** Primers, cycling conditions used and base-pairs (bp) amplified.

| Reference | Primer | Primer Direction (5' – 3') | PCR thermal cycling conditions | bp |
|---|---|---|---|---|
| 54 | LCO1490 | (F) GGTCAACAAATCATAAAGATATTGG | 1) 94°C (1 min); 2) 5 cycles: 94°C (30 s), 45°C (1 min 30 s), 72°C (1 min); 3) 35 cycles: 94°C (30 s), 51°C (1 min 30 s), 72°C (1 min); 4) 72°C (5 min). | 658 |
| | HCO2198 | (R) TAAACTTCAGGGTGACCAAAAAATCA | | |
| 56 | LoboF1 | (F) KBTCHACAAAYCAYAARGAYATHGG | 1) 94°C (2 min); 2) 35 cycles: 94°C (30 s), 46°C (1 min), 72°C (1 min); 3) 72°C (5 min). | 550 |
| | ArR5 | (R) GTRATIGCICCIGCIARIACIGG | | |
| 55 | LoboF1 | (F) KBTCHACAAAYCAYAARGAYATHGG | 1) 94°C (1 min); 2) 5 cycles: 94°C (30 s), 45°C (1 min 30 s), 72°C (1 min); 3) 45 cycles: 94°C (30 s), 54°C (1 min 30 s), 72°C (1 min); 4) 72°C (5 min). | 658 |
| | LoboR1 | (R) TAAACYTCWGGRTGWCCRAARAAYCA | | |

**Table S3.** List of peracaridean species used to complement the Bayesian clade credibility trees, with respective source, sampling location and number of individuals used.

| Order | Species | Source | Country/Island | Site | N | Latitude | Longitude |
|---|---|---|---|---|---|---|---|
| **AMPHIPODA** | *Ampithoe rubricata* | This study | Portugal | Dona Ana | 1 | 37.086969 | -8.667716 |
| **AMPHIPODA** | *Ampithoe sp.* | This study | Gran Canaria | Playa Melenara | 1 | 27.988891 | -15.370485 |
| **AMPHIPODA** | *Apohyale media* | 76 | Gran Canaria | Bañaderos | 1 | 28.149658 | -15.54018 |
| **AMPHIPODA** | *Apohyale prevostii* | 76 | Portugal | São Pedro Moel | 1 | 39.758015 | -9.033165 |
| **AMPHIPODA** | *Caprella liparotensis* | This study | Portugal | Dona Ana | 1 | 37.086969 | -8.667716 |
| **AMPHIPODA** | *Caprella mutica* | 77 | North Sea | Helgoland | 1 | 54.171000 | 7.889000 |
| **AMPHIPODA** | *Caprella penantis* | 55 | Portugal | Alentejo | 1 | 37.910000 | -8.800000 |
| **AMPHIPODA** | *Elasmopus canarius* | This study | Gran Canaria | Bañaderos | 2 | 28.149658 | -15.540180 |
| **AMPHIPODA** | *Elasmopus canarius* | This study | La Palma | El Faro | 1 | 28.457545 | -17.850340 |
| **AMPHIPODA** | *Elasmopus canarius* | This study | El Hierro | Arenas Blancas | 1 | 27.767189 | -18.121308 |
| **AMPHIPODA** | *Elasmopus rapax* | This study | Spain | Pedreira | 1 | 43.556170 | -8.274942 |
| **AMPHIPODA** | *Elasmopus vachoni* | This study | La Palma | La Fajana | 1 | 28.842276 | -17.794324 |
| **AMPHIPODA** | *Elasmopus vachoni* | This study | São Miguel | Ribeira Chã | 2 | 37.715417 | -25.486836 |
| **AMPHIPODA** | *Elasmopus vachoni* | This study | Santa Maria | São Lourenço | 1 | 36.988172 | -25.054211 |
| **AMPHIPODA** | *Hyale pontica* | 76 | Spain | Muxía | 1 | 43.092831 | -9.223431 |
| **AMPHIPODA** | *Hyalinae* | This study | Morocco | El Jadida | 1 | 33.264036 | -8.510717 |
| **AMPHIPODA** | *Jassa falcata* | This study | Spain | Pedreira | 1 | 43.55617 | -8.274942 |
| **AMPHIPODA** | *Jassa marmorata* | 77 | North Sea | Helgoland | 1 | 54.171000 | 7.889000 |
| **AMPHIPODA** | *Jassa ocia* | This study | Portugal | Ingrina | 1 | 37.045257 | -8.878047 |
| **AMPHIPODA** | *Jassa pusilla* | 77 | North Sea | Helgoland | 1 | 54.171000 | 7.889000 |
| **AMPHIPODA** | *Stenothoe marina* | 77 | North Sea | Helgoland | 1 | 54.171000 | 7.889000 |
| **ISOPODA** | *Campecopea hirsuta* | This study | Portugal | Ingrina | 1 | 37.045257 | -8.878047 |
| **ISOPODA** | *Cyathura carinata* | This study | Portugal | Viana do Castelo | 1 | 41.689194 | -8.847870 |
| **ISOPODA** | *Dynamene bidentata* | 78 | Gran Canaria | Bañaderos | 1 | 28.149658 | -15.540180 |
| **ISOPODA** | *Dynamene bidentata* | 78 | Morocco | El Jadida | 1 | 33.264036 | -8.510717 |
| **ISOPODA** | *Dynamene bidentata* | 78 | Portugal | Buarcos | 2 | 40.175976 | -8.900572 |
| **ISOPODA** | *Dynamene bidentata* | 78 | Portugal | Sines | 1 | 37.960884 | -8.887296 |
| **ISOPODA** | *Dynamene magnitorata* | 78 | La Palma | El Faro | 1 | 28.457545 | -17.85034 |
| **ISOPODA** | *Dynamene magnitorata* | 78 | São Miguel | Ribeira Chã | 1 | 37.715417 | -25.486836 |
| **ISOPODA** | *Dynamene magnitorata* | 78 | São Miguel | Ribeira Chã | 1 | 37.715417 | -25.486836 |
| **ISOPODA** | *Dynamene magnitorata* | 78 | Santa Maria | Praia Formosa | 1 | 36.949917 | -25.094989 |
| **ISOPODA** | *Dynamene magnitorata* | 78 | Portugal | Arrifes | 3 | 37.076052 | -8.276780 |
| **TANAIDACEA** | *Apseudes talpa* | 79 | Portugal | Mindelo | 1 | - | - |
| **TANAIDACEA** | *Tanais sp1* | This study | Santa Maria | Praia Formosa | 1 | 36.949917 | -25.094989 |
| **TANAIDACEA** | *Tanais sp2* | This study | Gran Canaria | Playa Melenara | 1 | 27.988891 | -15.370485 |
| **TANAIDACEA** | *Tanais sp3* | This study | Selvagens | Selvagem Pequena | 1 | 30.033233 | -16.016675 |
| **TANAIDACEA** | *Zeuxo exsargasso* | This study | Tenerife | Mal Paso | 1 | 28.416147 | -16.298656 |
| **TANAIDACEA** | *Zeuxo exsargasso* | This study | Porto Santo | Porto dos Frades | 1 | 33.072575 | -16.295666 |

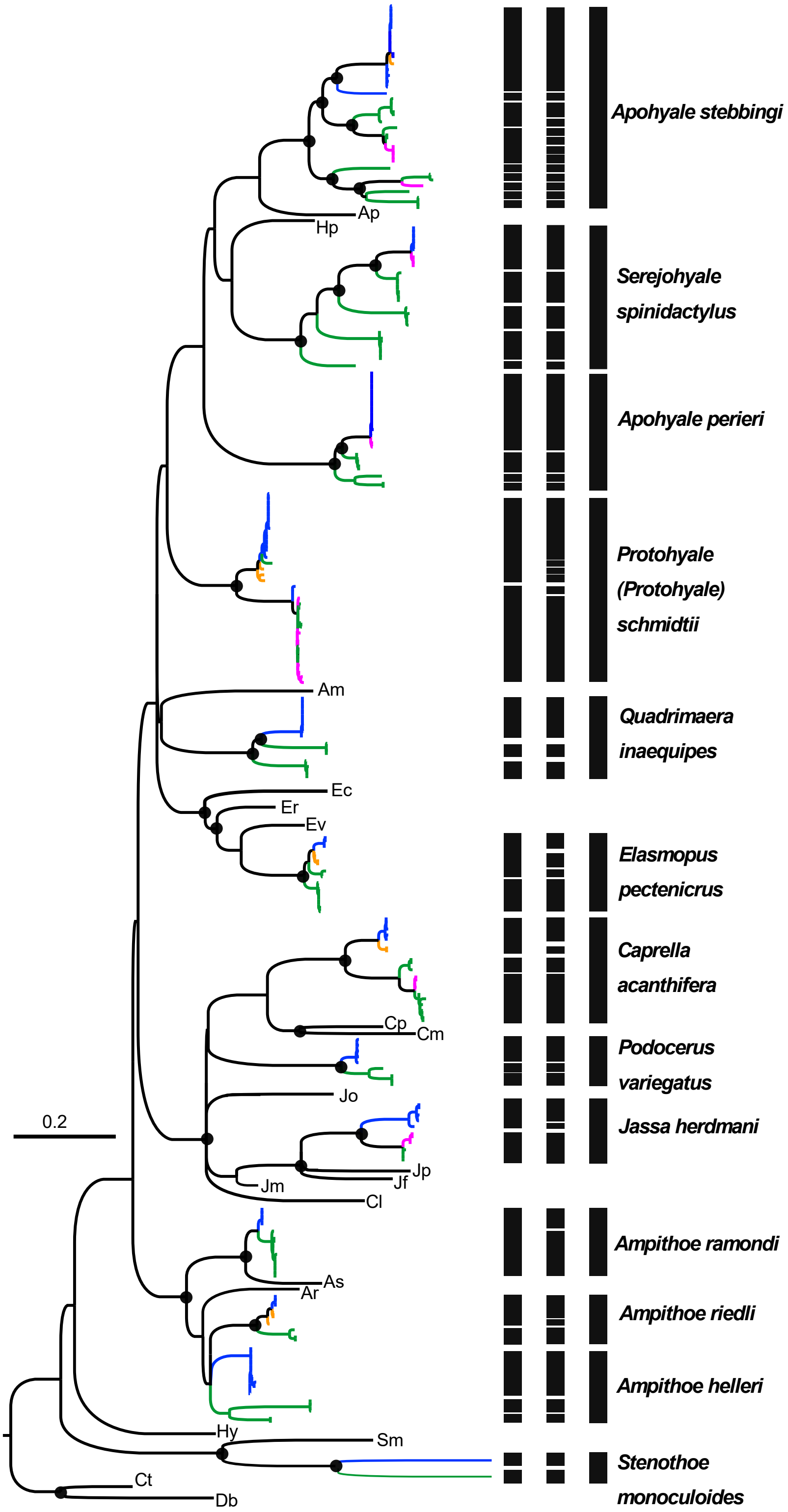


**Figure S1.** Bayesian clade credibility tree based on COI sequences of the amphipod species used in this study. Dotes (●) associated with nodes represent posterior probabilities higher than 0.90. Vertical black bars correspond to MOTUs obtained by the different methods of species delimitation applied (Table 2): A – lowest number of MOTUs, B – maximum number of MOTUs, C – morphospecies. *Cymodoce truncata* (Ct) and *Dynamene bidentata* (Db) were used as outgroup.

Lineages colors according with region: purple – AZ, green-MACA, orange – MORO, blue-IP.

Ap - *Apohyale prevostii*; Hp – *Hyale pontica*; Am – *Apohyale media*; Ec – *Elasmopus canarius*; Er – *Elasmopus rapax*; Ev – *Elasmopus vachoni*; Cp – *Caprella penantis*; Cp – *Caprella mutica*; Jf – *Jassa falcata*; Jo – *Jassa ocia*; Jp – *Jassa pusilla*; Jm – *Jassa marmorata*; Cl – *Capella liparotensis*; As – *Ampithoe sp.*; Ar – *Ampithoe rubricata*; Hy – Hyalinae; Sm – *Stenothoe marina*.

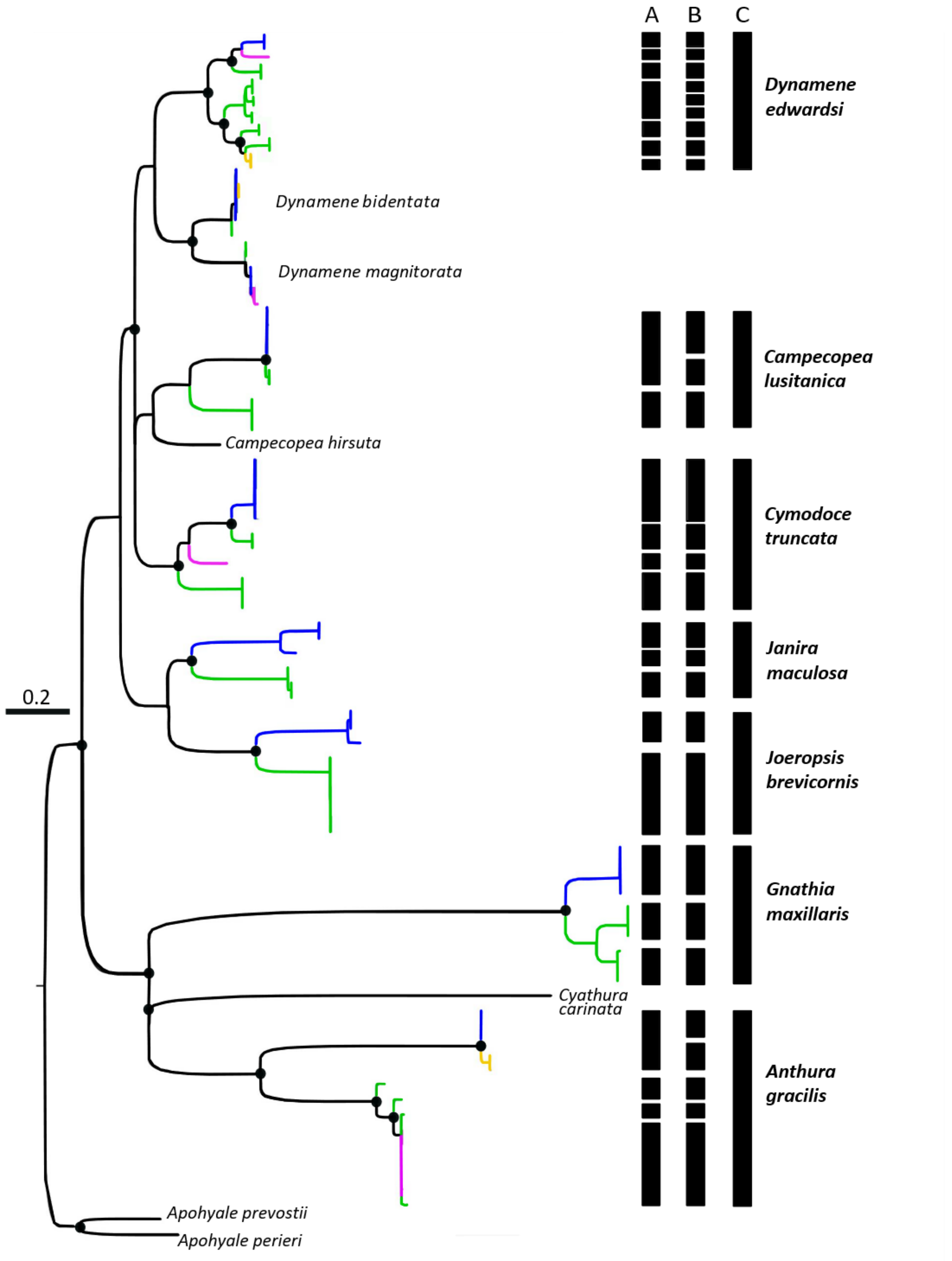


**Figure S2.** Bayesian clade credibility tree based on COI sequences of the isopod species used in this study. Dotes (●) associated with nodes represent posterior probabilities higher than 0.90. Vertical black bars correspond to MOTUs obtained by the different methods of species delimitation applied (Table 2): A – lowest number of MOTUs, B – maximum number of MOTUs, C – morphospecies. *Apohyale prevostii* and *Apohyale perieri* were used as outgroup.

Lineages colors according with region: purple – AZ, green-MACA, orange – MORO, blue-IP.

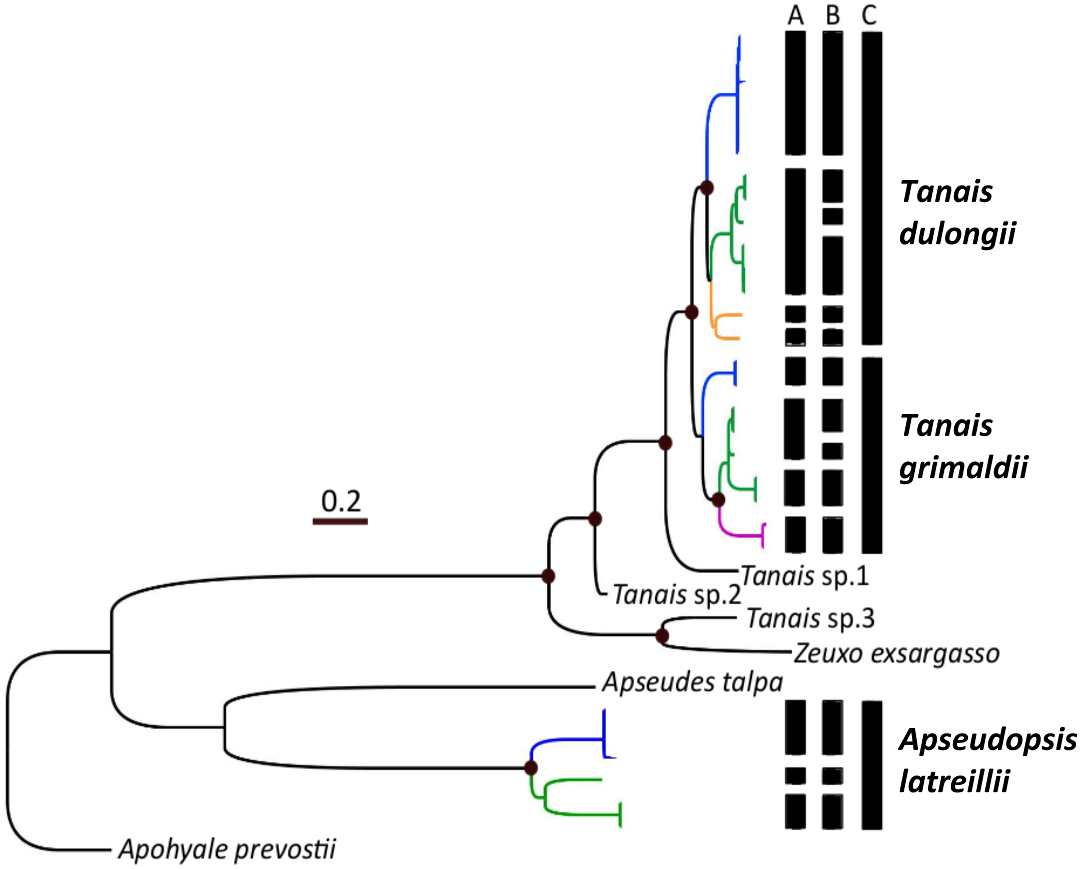


**Figure S3.** Bayesian clade credibility tree based on COI sequences of the tanaid species used in this study. Dotes (●) associated with nodes represent posterior probabilities higher than 0.90. Vertical black bars correspond to MOTUs obtained by the different methods of species delimitation applied (Table 2): A – lowest number of MOTUs, B – maximum number of MOTUs, C – morphospecies. *Apohyale prevostii* was used as outgroup. Lineages colours according with region: purple – AZ, green-MACA, orange – MORO, blue-IP.

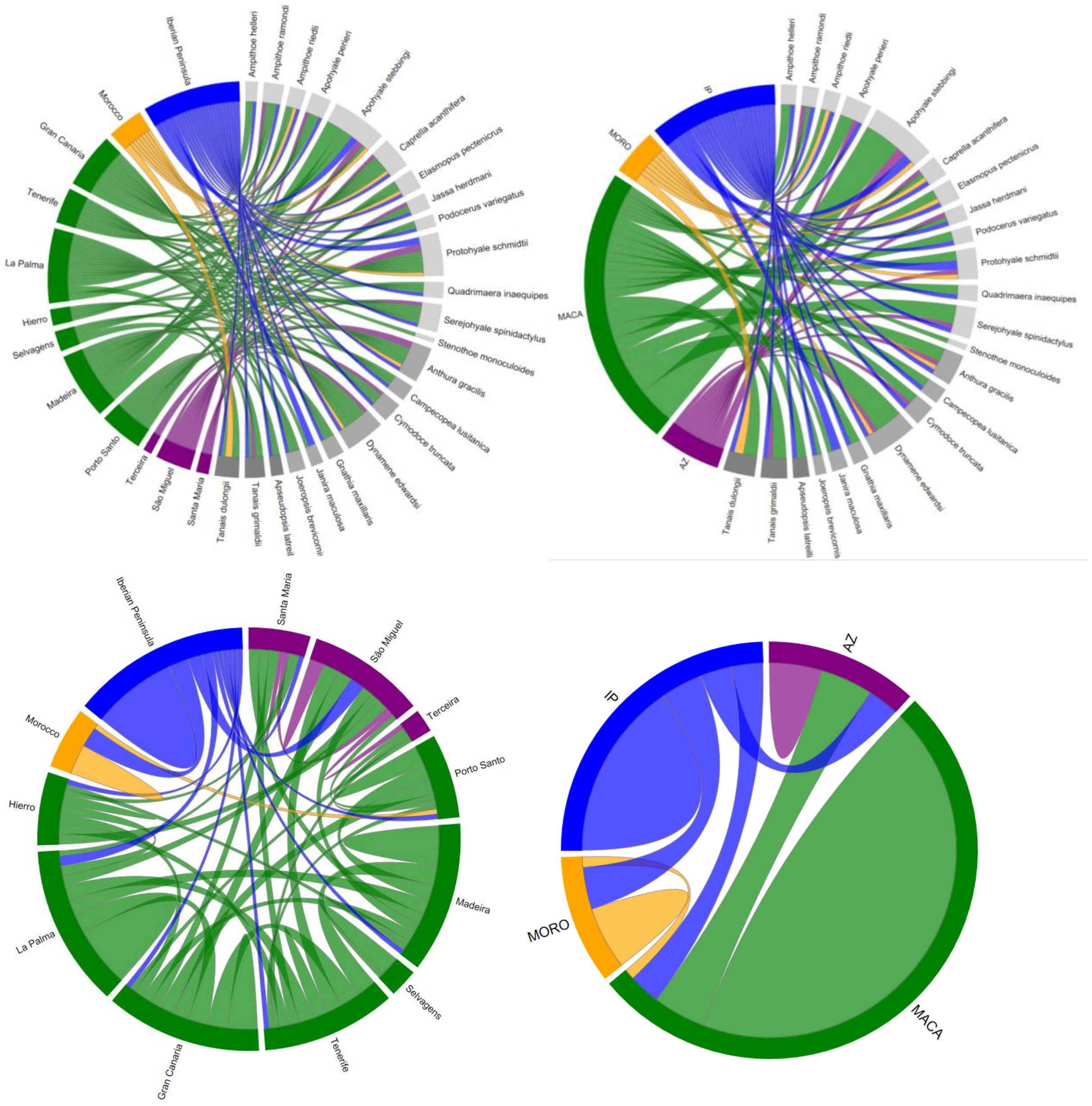


**Figure S4.** Upper: MOTUs present in each location (left) and region (right) for each morphospecies. Lower: Private and shared MOTUs between locations (left) and regions (right).

For interactive diagrams, see respectively: upper left - http://rpubs.com/Vieira/Diagram; upper right - http://rpubs.com/Vieira/Diagram2; lower left - http://rpubs.com/Vieira/Diagram3 and lower right - http://rpubs.com/Vieira/Diagram4. Colors according with region: purple - AZ, green - MACA, orange - MORO, blue - IP and orders: lightgrey - Amphipoda, grey - Isopoda, darkgrey - Tanaidacea.

An interactive app created with the package ‘shiny’ [80] in R 3.5.0 [2] can be consulted at https://pedroevieira.shinyapps.io/test/ to visualize the four diagrams.

### References


1. Cheng J, Karambelkar B, Xie Y. 2018 leaflet: Create Interactive Web Maps with the JavaScript 'Leaflet' Library. R package version 2.0.1. https://CRAN.R-project.org/package=leaflet.
2. R Core Team. 2018 R: A language and environment for statistical computing. R Foundation for Statistical Computing, Vienna, Austria. https://www.R-project.org/.
3. Chevreux E, Fage L. 1925 Faune de France. 9, Amphipodes. Lechevalier, Paris (France).
4. Naylor E. 1972 British Marine Isopods. Keys and Notes for the Identification of the Species– Synopses of the British Fauna. Academc Press, London-New York, 90 pp.
5. Lincoln R. 1979 British marine amphipoda: Gammaridea. British Museum (Natural History), London (United Kigdom).
6. Ruffo S. 1983 The Amphipoda of the Mediterranean. Memoires de l'Institut Oceanographique, Monaco.
7. Holdich DM, Jones J. 1983 Tanaids. Keys and notes for the identification of the species, 27. The Linnean Society of London, London (Unites Kigdom).
8. Harrison K, Ellis JP. 1991 The genera of the Saphaeromatidae (Crustacea: Isopoda): a key and distribution list. Invertebr. Syst. **5**, 915–952.
9. Hayward P, Ryland J. 1995 Handbook of the marine fauna of North-west Europe. Oxford University press, Oxford (United Kingdom), 816 pp.
10. Dallwitz M, Paine T, Zurcher E. 2000 Principles of interactive keys: onwards. Available from: http://delta-intkey.com.
11. Lowry J, Springthorpe R. 2001 Amphipoda: Families. Available from: http://www.crustacea.net/.
12. Larsen K, Larsen K. 2002 Tanaidacea: Families. Available from: http://crustacea.net.
13. Keable S, Poore G, Wilson G. 2002 Australian Isopoda: Families. Available from: http://crustacea.net.
14. Coleman CO, Lowry J, Macfarlane T. 2010 DELTA for Beginners. An introduction into the taxonomy software package DELTA. ZooKeys **45**, 1–75.
15. Hispano C, Bultó P, Blanch AR. 2014 Life cycle of the fish parasite Gnathia maxillaris (Crustacea: Isopoda: Gnathiidae). Folia Parasitol. (Praha) **61**, 277–284.
16. Holdich DM. 1968 A systematic revision of the genus *Dynamene* Crustacea Isopoda with a description of three new species. Pubbl. Stn. zool. Napoli **36**, 401–426.
17. Vieira PE, Queiroga H, Costa FO, Holdich DM. 2016 Distribution and species identification in the crustacean isopod genus *Dynamene* Leach, 1814 along the North East Atlantic-Black Sea axis. ZooKeys **635**, 1–29. (doi: 10.3897/zookeys.635.10240).
18. Nolting C, Reboreda P, Wägele JW. 1998 Systematic Revision of the Genus *Anoplocopea* Racovitza, 1907 (Crustacea: Isopoda) with a Description of a New Species from the Atlantic Coast of the Iberian Peninsula. Zoosystematics Evol. **74**, 19–41.
19. Bruce NL, Holdich DM. 2002 Revision of the isopod crustacean genus *Campecopea* (Flabellifera: Sphaeromatidae) with discussion of the phylogenetic significance of dorsal processes. J. Mar. Biol. Assoc. U. K. **82**, 51–68.
20. Khalaji-Pirbalouty V, Bruce NL, Wägele J.W. 2013 The genus *Cymodoce* Leach, 1814 (Crustacea: Isopoda: Sphaeromatidae) in the Persian Gulf with description of a new species. Zootaxa **3686**, 501–533.
21. Khalaji-Pirbalouty V, Raupach MJ. 2014 A new species of *Cymodoce* Leach, 1814 (Crustacea: Isopoda: Sphaeromatidae) based on morphological and molecular data, with a key to the Northern Indian Ocean species. Zootaxa **3826**, 230. (doi: 10.11646/zootaxa.3826.1.7).
22. Poore G. 2001 Families and genera of Isopoda Anthuridea. In: Kensley B, Brusca RC (eds) Isopod systematics and evolution. Balkema Publications, Rotterdam, pp 63–173.
23. Esquete P, Bamber RN, Moreira J, Troncoso JS. 2012 Redescription and postmarsupial development of *Apseudopsis latreillii* (Crustacea: Tanaidacea). J. Mar. Biol. Assoc. U. K. **92**, 1023–1041. (doi: 10.1017/S0025315411002086).
24. Esquete P, Ramos E, Riera R. 2016 New data on the Tanaidacea (Crustacea: Peracarida) from the Canary Islands, with a description of a new species of *Apseudopsis*. Zootaxa **4093**, 248–260. (doi: 10.11646/zootaxa.4093.2.6).
25. Bamber RN. 2012 Littoral Tanaidacea (Crustacea: Peracarida) from Macaronesia: allopatry and provenance in recent habitats. J. Mar. Biol. Assoc. U. K. **92**, 1095–1116. (doi: 10.1017/S0025315412000252).
26. Bamber RN, Robbins R. 2009 The soft-sediment infauna off São Miguel, Azores, and a comparison with other Azorean invertebrate habitats. Açoreana **6**, 201–210.
27. Riera R, Guerra-García J, Brito M, Núñez J. 2003 Estudio de los caprélidos de Lanzarote, islas Canarias (Crustacea: Amphipoda: Caprellidea). Vieraea **31**, 157–166.

28. Lacerda MB, Masunari S. 2011 Chave de identificação para caprelídeos (Crustacea, Amphipoda) do litoral dos Estados do Paraná e de Santa Catarina. Biota Neotropica **11**, 379–390.
29. Guerra-García JM, Ros M, Izquierdo D, Soler-Hurtado MM. 2012 The invasive *Asparagopsis armata* versus the native *Corallina elongata*: Differences in associated peracarid assemblages. J. Exp. Mar. Biol. Ecol. **416–417**, 121–128. (doi: 10.1017/S0025315411002086).
30. Guerra-García JM. et al. 2013 An illustrated key to the soft-bottom caprellids (Crustacea: Amphipoda) of the Iberian Peninsula and remarks to their ecological distribution along the Andalusian coast. Helgol. Mar. Res. **67**, 321–336. (doi: 10.1007/s10152-012-0324-1).
31. Krapp T, Rampin M, Libertini A. 2008 A cytogenetical study of Ischyroceridae (Amphipoda) allows the identification of a new species, *Jassa cadetta* sp. n., in the Lagoon of Venice. Org. Divers. Evol. **8**, 337–345.
32. Conlan KE. 1990 Revision of the crustacean amphipod genus *Jassa* Leach (Corophioidea: Ischyroceridae). Can. J. Zool. **68**, 2031–2075.
33. Lowry JK, Hughes LE. 2009 Maeridae, the *Elasmopus* group. Zootaxa **2260**, 643–702.
34. Vader W, Krapp-Schickel T. 2012 On some maerid and melitid material (Crustacea: Amphipoda) collected by the Hourglass Cruises (Florida). Part 2: Genera *Dulichiella* and *Elasmopus*, with a key to world *Elasmopus*. J. Nat. Hist. **46**, 1179–1218.
35. Gouillieux B, Sorbe JC. 2015 *Elasmopus thalyae* sp. nov. (Crustacea: Amphipoda: Maeridae), a new benthic species from soft and hard bottoms of Arcachon Bay (SE Bay of Biscay). Zootaxa **3905**, 107–118. (doi: 10.11646/zootaxa.3905.1.6).
36. Alves J, Johnsson R, Senna AR. 2016 On the genus *Elasmopus* Costa, 1853 from the Northeastern Coast of Brazil with five new species and new records. Zootaxa **4184**, 1–40. (doi: 10.11646/zootaxa.4184.1.1).
37. Krapp-Schickel T. 2006 New Australian stenothoids (Crustacea, Amphipoda) with key to all *Stenothoe* species. Boll. Mus. Civ. Storia Nat. Verona Bot. Zool. **30**, 39–56.
38. Krapp-Schickel T. 2015 Minute but constant morphological differences within members of Stenothoidae: the *Stenothoe gallensis* group with four new members, keys to *Stenothoe* worldwide, a new species of *Parametopa* and *Sudanea* n. gen. (Crustacea: Amphipoda). J. Nat. Hist. **49**, 2309–2377. (doi: 10.1080/00222933.2015.1021873).
39. Conlan KE. 1982 Revision of the gammaridean amphipod family Ampithoidae using numerical analytical methods. Can. J. Zool. **60**, 2015–2027.
40. Hughes L, Kilgallen N, Lowry J, Peart R. 2008 Ampithoidae (Amphipoda): World Genera and Australian, Northeast Atlantic and Mediterranean Species. Available from: http://crustacea.net.
41. Krapp-Schickel T, Ruffo S. 1990 Marine Amphipods of the Canary Islands with description of a new species of *Elasmopus*. Miscellània Zoològica **14**, 53–58.
42. Castelló J, Carballo JL. 2001 Isopod fauna, excluding epicaridea, from the strait of Gibraltar and nearby areas (Southern Iberian Peninsula). Sci. Mar. **65**, 221–241.
43. Junoy J, Castelló-Escandell J. 2003 Catálogo de las especies ibéricas y baleares de isópodos marinos (Crustacea: Isopoda). Bol. Inst. Esp. Oceanogr. **19**, 293-326.
44. Costello M, Emblow C, White R. 2001 European Register of Marine Species. A check-list of the marine species in Europe and a bibliography of guides to their identification. Patrimoines naturels. **50**, 463.
45. Gomes S, Lima F, Queiroz N, Ribeiro P, Santos A. 2006 Biogeographic Patterns of Intertidal Macroinvertebrates and their Association with Macroalgae Distribution along the Portuguese Coast. Hydrobiologia **555**, 185–192.
46. Boyko C et al. 2008 World Marine, Freshwater and Terrestrial Isopod Crustaceans database. Available from: http://www.marinespecies.org/isopoda.
47. Borges P et al. 2010 A list of the terrestrial and marine biota from the Azores, 1st edition. Princípia Editora, Cascais (Portugal) 429pp.
48. Izquierdo D, Guerra-García JM. 2011 Distribution patterns of the peracarid crustaceans associated with the alga *Corallina elongata* along the intertidal rocky shores of the Iberian Peninsula. Helgol. Mar. Res. **65**, 233–243. (doi: 10.1007/s10152-010-0219-y).
49. Anderson G. 2016 Tanaidacea-Thirty Years of Scholarship, Version 2.0. http://aquila.usm.edu/tanaids30.
50. Horton T et al. 2017 World Register of Marine Species (WoRMS). Available from: http://www.marinespecies.org (January 26, 2017).
51. Horton, T. et al. 2017 World Amphipoda Database. Available from: http://www.marinespecies.org/amphipoda (January 26, 2017).
52. Castelló J, Junoy J. 2007 Catálogo de las especies de isópodos marinos (Crustacea : Isopoda) de los archipiélagos macaronésicos. Bol. Inst. Esp. Oceanogr. **23**.
53. Guerra-García JM, Baeza-Rojano E, Cabezas MP, García-Gómez JC. 2011 Vertical distribution and seasonality of peracarid crustaceans associated with intertidal macroalgae. J. Sea Res. **65**, 256–264.

54. Folmer O, Black B, Hoeh W, Lutz R, Vrijenhoek R. 1994 DNA primers for amplification of mitochondrial cytochrome c oxidase subunit I from diverse metazoan invertebrates. Mol. Mar. Biol. Biotechnol. **3**, 294–299.
55. Lobo J et al. 2013 Enhanced primers for amplification of DNA barcodes from a broad range of marine metazoans. BMC Ecol. **13**, 34. (doi: 10.1186/1472-6785-13-34).
56. Gibson J et al. 2014 Simultaneous assessment of the macrobiome and microbiome in a bulk sample of tropical arthropods through DNA metasystematics. Proc. Natl. Acad. Sci. **111**, 8007–8012. (doi: 10.1073/pnas.1406468111).
57. Thompson JD, Higgins DG, Gibson TJ. 1994 CLUSTAL W: improving the sensitivity of progressive multiple sequence alignment through sequence weighting, position-specific gap penalties and weight matrix choice. Nucleic Acids Res. **22**, 4673–4680.
58. Kumar S, Stecher G, Tamura K. 2016 MEGA7: Molecular Evolutionary Genetics Analysis Version 7.0 for Bigger Datasets. Mol. Biol. Evol. **33**, 1870–1874. (doi: 10.1093/molbev/msw054).
59. Librado P, Rozas J. 2009 DnaSP v5: a software for comprehensive analysis of DNA polymorphism data. Bioinforma. Oxf. Engl. **25**, 1451–1452. (doi: 10.1093/bioinformatics/btp187).
60. Altschul SF, Gish W, Miller W, Myers EW, Lipman DJ. 1990 Basic local alignment search tool. J. Mol. Biol. **215**, 403–410.
61. Ratnasingham S, Hebert PDN. 2007 bold: The Barcode of Life Data System (http://www.barcodinglife.org). Mol. Ecol. Notes **7**, 355–364. (doi: 10.1111/j.1471-8286.2007.01678.x).
62. Ronquist F et al. 2012 MrBayes 3.2: efficient Bayesian phylogenetic inference and model choice across a large model space. Syst. Biol. **61**, 539–542. (doi: 10.1093/sysbio/sys029).
63. Puillandre N, Lambert A, Brouillet S, Achaz G. 2012 ABGD, Automatic Barcode Gap Discovery for primary species delimitation. Mol. Ecol. **21**, 1864–1877. (doi: 10.1111/j.1365-294X.2011.05239.x).
64. Ratnasingham S, Hebert PDN. 2013 A DNA-Based Registry for All Animal Species: The Barcode Index Number (BIN) System. PLOS ONE **8**, e66213. (doi: 10.1371/journal.pone.0066213).
65. Zhang J, Kapli P, Pavlidis P, Stamatakis A. 2013 A general species delimitation method with applications to phylogenetic placements. Bioinforma. Oxf. Engl. **29**, 2869–2876. (doi: 0.1093/bioinformatics/btt499).
66. Fujisawa T, Barraclough TG. 2013 Delimiting species using single-locus data and the Generalized Mixed Yule Coalescent approach: a revised method and evaluation on simulated data sets. Syst. Biol. **62**, 707–724. (doi: 10.1093/sysbio/syt033).
67. Pons J et al. 2006 Sequence-based species delimitation for the DNA taxonomy of undescribed insects. Syst. Biol. **55**, 595–609.
68. Monaghan MT et al. 2009 Accelerated species inventory on Madagascar using coalescent-based models of species delineation. Syst. Biol. **58**, 298–311.
69. Bouckaert R et al. 2014 BEAST 2: A Software Platform for Bayesian Evolutionary Analysis. PLOS Comput. Biol. **10**, e1003537. (doi: 10.1371/journal.pcbi.1003537).
70. Rambaut A, Suchard M, Xie D, Drummond AJ. 2014 Tracer v1.6. Available from http://beast.bio.ed.ac.uk/Tracer.
71. Ezard T, Fujisawa T, Barraclough TG. 2009 SPLITS: species' limits by threshold statistics. R package version 1.0–18/r45. Available at: http://R-Forge.R-project.org/projects/splits/.
72. Clement M, Posada D, Crandall KA. 2000 TCS: a computer program to estimate gene genealogies. Mol. Ecol. **9**, 1657–1659.
73. Sauer J, Hausdorf B. 2012 A comparison of DNA-based methods for delimiting species in a Cretan land snail radiation reveals shortcomings of exclusively molecular taxonomy. Cladistics **28**, 300–316.
74. Yang Z, Algesheimer R, Tessone CJ. 2016 A Comparative Analysis of Community Detection Algorithms on Artificial Networks. Sci. Rep. **6**, 30750. (doi: 10.1038/srep30750).
75. Lobo J et al. 2016 Contrasting morphological and DNA barcode-suggested species boundaries among shallow-water amphipod fauna from the southern European Atlantic coast. Genome. **60,** 147-157. (doi: 10.1139/gen-2016-0009).
76. Desiderato A et al. 2019 Macaronesian islands as promoters of diversification in amphipods: The remarkable case of the family Hyalidae (Crustacea, Amphipoda). Zool. Scr. **48**, 359–375. (doi: 10.1111/zsc.12339).
77. Raupach MJ et al. 2015 The Application of DNA Barcodes for the Identification of Marine Crustaceans from the North Sea and Adjacent Regions. PLOS ONE **10**, e0139421. (doi: 10.1371/journal.pone.0139421).
78. Vieira PE et al. 2019 Deep segregation in the open ocean: Macaronesia as an evolutionary hotspot for low dispersal marine invertebrates. Mol. Ecol. **28**, 1784–1800. (doi: 10.1111/mec.15052).
79. Larsen K, Bertocci I, Froufe E. 2011 *Apseudes talpa* revisited (Crustacea; Tanaidacea). The impact on apseudidaen systematics. Zootaxa **2886**, 19–30. (doi: 10.11646/zootaxa.2886.1.2).

80. Chang W, Cheng J, Allaire J, Xie Y, McPherson J. 2018 shiny: Web Application Framework for R. R package version 1.1.0. https://CRAN.R-project.org/package=shiny.
